# Spatially-distinct programming of macrophage diversity within the granulomas of *Mycobacterium tuberculosis* infected nonhuman primates

**DOI:** 10.1101/2025.06.12.659348

**Authors:** Davide Pisu, Persis S. Sunny, Molly L. Nelson, Beth F. Junecko, Roopa Venugopalan, Mark A. Rodgers, H. Jacob Borish, Pauline Maiello, Charles A. Scanga, Philana Ling Lin, JoAnne L. Flynn, David G. Russell, Joshua T. Mattila

## Abstract

Tuberculosis (TB), caused by *Mycobacterium tuberculosis* (Mtb), is defined by granulomas— immune aggregates that either contain or support bacterial replication. Macrophages, fundamental components of these lesions, are crucial to TB pathogenesis, yet their phenotypic and functional diversity is incompletely understood. Here, we used single-cell RNA sequencing and immunofluorescence to profile macrophages in lung tissue and granulomas from a nonhuman primate model of early TB. We identified distinct subsets, including embryonic-origin tissue-resident alveolar macrophages and monocyte-derived alveolar and interstitial macrophages, with distinct spatial localization in granulomas. Tissue-resident alveolar macrophages and a subset undergoing epithelial-to-mesenchymal transition accounted for the highest frequency of Mtb-infected cells. Infected cells exhibited differential expression of immune- and migration-associated genes compared to uninfected counterparts, suggesting Mtb either induces or exploits these pathways as a survival strategy. These findings highlight macrophage heterogeneity as a major driver of differential susceptibility to Mtb and provide insights relevant to future immunomodulatory strategies.

## Introduction

Tuberculosis (TB) remains a major global health challenge, causing over 1.3 million deaths annually despite continued efforts to improve treatment and prevention strategies[1, 2]. TB begins when *Mycobacterium tuberculosis* (Mtb) is inhaled into the lungs, initiating immune responses that culminate in the formation of granulomas—complex immune cell aggregates that are the hallmark of the disease. These lesions can effectively limit bacterial replication or, paradoxically, serve as a reservoir for Mtb growth and persistence[3–6]. Macrophages are central to granuloma biology, acting as intracellular niches for Mtb, antimicrobial effectors that limit bacterial replication, and key regulators of the immune networks shaping granuloma functionality[7–10]. However, the phenotypic and functional diversity of macrophage subsets within granulomas remains incompletely understood, and expanding this knowledge is critical for refining immunological correlates of protection and developing targeted TB interventions.

Multi-modal single-cell RNA sequencing (scRNA-seq) has revolutionized our ability to dissect complex population structures and cell-cell interactions within TB-affected tissues[7, 8, 11–14]. This technology has provided detailed insights into the immune cell composition of Mtb-infected nonhuman primate (NHP) lungs[7], detailed the functional dynamics of granulomas following CD4 and CD8 T cell depletion[11, 15], and linked immune cell subsets to bacterial burden[8]. Although these studies revealed the presence of major diverse myeloid subsets in Mtb-infected NHP lungs, shedding light on pathways that influence macrophage differentiation[16], they did not focus specifically on identifying macrophage subpopulations or their unique contributions to granuloma organization and host defense. Consequently, gaps remain in our understanding of which macrophage subsets facilitate or restrict Mtb growth, and how these subsets shape protective versus pathogenic outcomes.

Building on approaches used in murine models—which employed fluorescent fitness reporter strains to profile both host and pathogen phenotypes with high resolution, revealing how specific macrophage subsets are associated with distinct bacterial states and fitness levels [12, 13, 17–19]—we adapted a similar strategy for the NHP model of TB, where disease pathology more closely parallels human infection.

By using a virulent Mtb strain expressing the fluorescent protein mCherry and integrating immunostaining, flow cytometry, and multi-modal scRNA-seq analyses, we provide an in-depth understanding of macrophage phenotypes in early TB in NHP lungs by detailing the composition, spatial localization, and infection status of alveolar (AMs) and interstitial macrophages (IMs). We demonstrate that extravasating monocytes give rise to diverse macrophage subsets that drive the local immune response. Through the identification of novel subset-specific markers, we show that AM and IM subpopulations occupy distinct niches within caseous granulomas, and that monocyte-derived AMs (moAMs) exhibit pronounced inflammatory phenotypes in these lesions. Although multiple macrophage subsets harbor Mtb, we discovered a minor mesenchymal-like macrophage population that is disproportionately infected, identifying an opportunity induced or exploited by Mtb to subvert host immunity. Altogether, these findings provide a novel perspective on macrophage diversity and dynamics in the early stages of TB—an interval critical for shaping long-term disease outcomes[8]. By identifying key macrophage subsets and their interactions with Mtb, we provide a novel framework for targeting macrophage-specific pathways in granulomas with implications for improving therapeutic strategies for TB.

## Materials and Methods

### Animal ethics statement

All experimental manipulations, procedures, protocols, and husbandry for the non-human primate studies were approved by the University of Pittsburgh School of Medicine Institutional Animal Care and Use Committee (IACUC) under protocol number 20067350. The University of Pittsburgh’s IACUC adheres to national guidelines established in the Animal Welfare Act (7 U.S.C. Sections 2131–2159) and the Guide for the Care and Use of Laboratory Animals (eighth edition), as mandated by the U.S. Public Health Service Policy.

### Mtb Strains

The parental strain employed for all experiments was *Mycobacterium tuberculosis* Erdman (ATCC 35801). Fluorescent reporter strains, including smyc′::mCherry and smyc′::mCherry/hspx’::GFP, were previously described [18, 20]. Mtb strains were cultivated at 37°C until they reached the mid-log phase in Middlebrook 7H9 broth enriched with 10% OADC (Becton, Dickinson and Company), 0.2% glycerol, and 0.05% tyloxapol (Sigma-Aldrich, St. Louis, MO). Hygromycin B (50 mg/ml) was used for selection of fluorescent strains.

Bacterial aliquots were prepared in 10% glycerol, titrated, and preserved at −80°C, following the protocol detailed in Pisu et al[21]. To select for bacterial colonies that retained high virulence traits and minimized the loss of the fluorescent protein in NHP infections, the bacterial aliquots were used to infect C57BL/6J mice according to previously-published protocols [21]. Three weeks post-infection, mice were sacrificed, and the lung lobes were homogenized in PBS containing 0.05% tyloxapol (Sigma-Aldrich) for plating on 7H10 agar plates supplemented with Hygromycin B (50 mg/ml). Mtb colonies showing the brightest expression of the mCherry signal following mouse infection were isolated and expanded in 7H9/OADC broth supplemented with Hygromycin B.

Aliquots from these highly expressing and virulent mCherry colonies were prepared in 10% glycerol, titrated, preserved at −80°C, and used for NHP infections. All bacterial aliquots used for NHP infections were tested for phthiocerol dimycocerosate (PDIM) expression and confirmed to be PDIM^+^.

### NHP infections, monitoring, and tissue sampling

Three Mauritian cynomolgus macaques (*Macaca fascicularis*, listed hereafter as animals 30320, 30420, and 30520) were purchased from Bioculture Mauritius and quarantined before being infected via bronchoscopic instillation of recombinant mCherry-expressing strains of Mtb that were based on the Erdman strain. Animals were infected bilaterally with 17-40 CFU (Supplementary Table 1) with the intent of generating sufficient levels of disease for sampling and scRNA-seq analysis. The animals were infected for 28-30 days before undergoing PET-CT imaging (Supplementary Table 2) with the glucose analog ^18^F-FDG [22, 23] to confirm infection and to identify granulomas to be harvested at necropsy. The overall infection duration was 32-33 days (Supplementary Table 1) before the animals were humanely euthanized and necropsied as previously described [24].

At necropsy, pieces of non-diseased lung were selected from lung lobes that lacked granuloma-like ^18^F-FDG retention by PET-CT and the absence of grossly visible tuberculous pathology was confirmed by visual inspection while the tissue was being harvested. Tissues meeting this criterion are described as non-diseased throughout this work although small numbers of bacteria were recovered from several pieces. Granuloma-containing diseased lung was identified by its radiographic appearance on CT, ^18^F-FDG retention, and the presence of grossly visible disease. After excision, the tissues were kept in cold RPMI until processed for freezing or plating. Random pieces of lung tissue from all animals were fixed with 10% neutral buffered formalin, embedded in paraffin, and cut into 5 μm sections for hematoxylin and eosin (H&E) staining for histologic examination. To further confirm infection, select samples of non-diseased lung and lymph nodes were mechanically disaggregated as previously described[24] and plated on 7H11 plates.

Granulomatous tissue was reserved for scRNA-seq analysis and was not sampled for bacterial burdens; thus, CFU per thoracic lymph node data are presented as representative data (Supplementary Table 3) to confirm Mtb infection and not as an overall quantification of bacterial burdens per animal.

Our rationale for using frozen lung tissue as a source of isolated cells for scRNA-seq was based on previously published literature [25–29] and validated to confirm this approach yielded viable macrophages that were amenable to flow cytometry (Supplementary Figure 2, panels A-D). Non-diseased tissue and granuloma-containing tissues were cut into pieces approximately 0.5 cm on a side and placed in a 1.5 mL cryovial that was filled with CellBanker 2 freezing medium (AMSBIO LLC, Cambridge, MA) so that one third of the tube was filled with tissue and the remaining volume was filled with CellBanker 2. The tubes were mixed by inversion to ensure penetrance of the freezing medium, placed with an isopropanol-containing Mr. Frosty cell freezing container (Thermo Fisher Scientific, Waltham, MA), and then stored in a -80°C freezer. The samples were shipped to Cornell University on dry ice to be processed for scRNA-seq.

### Lung cell isolation, staining and flow sorting

1.5mL cryovials containing non-diseased and granuloma-containing lung tissues were quickly thawed at 37°C for 2 minutes using a water bath. Upon thawing, the contents of each tube were poured over a 70μM cell strainer and into a 50mL conical tube. Granulomatous or non-diseased tissues from each animal were pooled together by disease state to maximize the number of total cells and infected cells recovered during sorting. Using forceps, multiple tissue pieces were combined into granuloma-only or non-diseased lung-only aliquots, with separate tubes per animal, and transferred into GentleMacs C tubes (max 5-6 tissue sections/C tube; Miltenyi Biotec, Gaithersburg, MD) prefilled with 5mL of enzyme solution from the Tumor Dissociation Kit (200uL enzyme H, 100uL enzyme R, 25uL enzyme A, 4.675 mL RPMI 1640; Miltenyi Biotec).

To minimize unwanted changes in the gene expression profile of isolated tissues, quick tissue homogenization was done by running a custom GentleMACS program for 15’ at 37°C (+180 rpm for 20”, -180 rpm for 20”, loop 22X). The downstream steps were performed at 4°C whenever possible. Lung cell suspensions obtained after running the GentleMACs program were then passed through a 70uM strainer, spun at 500g for 5’ at 4°C, incubated for 5’ in 1mL of ACK lysis buffer at room temperature (RT) (LONZA, Durham, NC), the reaction quenched with 10mL of cold PBS containing 5% FBS, spun down at 500g for 5’ at 4°C and finally resuspended in cold PBS containing 5% FBS.

The single cell suspensions were then stained with 5uL of a CD45 antibody (BD, cat #561294) for 20’ at 4°C, washed twice in cold PBS containing 5% FBS and centrifuged at 500g for 5’. The flow-stained lung suspensions were then resuspended in 200 µL of cell staining buffer containing 20 µL of Human TruStain FcX blocking reagent (BioLegend, San Diego, CA) and incubated for 10 min at 4°C. An antibody-derived tag (ADT) cocktail mix (200 µL) was then added to the samples, and the cells were further incubated for 30 min at 4°C. After two washes in 1mL of cell staining buffer, the flow- and ADT-stained cells were resuspended in sorting buffer (PBS, 1% FBS, 5 mM EDTA, and 25 mM HEPES), filtered through a 40µM strainer, and sorted. Throughout the sorting, samples were kept at 4°C and directly collected into Cell Staining Buffer (BioLegend).

### scRNA-seq libraries preparation and sequencing

Sorted cells were centrifuged at 500g for 5 min and resuspended in 1×Dulbecco’s PBS. The samples were fixed by slowly adding ice-cold methanol to a final concentration of 90% (vol/vol) and stored at −20°C overnight. Post-fixation, the samples were brought out of the BSL3 facility, equilibrated on ice for 15 min, and washed twice with rehydration buffer (1× Dulbecco’s PBS with 1.0% bovine serum albumin (BSA; Thermo Fisher Scientific) and 0.5 U/µl RNase Inhibitor (Sigma-Aldrich)[12, 21]. The cell count was determined prior to loading onto the 10X G chip for 3’ scRNA-seq (10X Genomics, Pleasanton, CA). For mRNA library preparation, we adapted the 10X protocol (CG000206 Rev D), making a minor alteration in step 2.2. Specifically, we incorporated 1 μl of ADT additive primer (0.2 μM stock) as previously described[30]. ADT libraries were prepared according to BioLegend’s standard protocols. The mRNA and ADT libraries underwent quality control assessment using an Agilent Fragment Analyzer (Agilent, Santa Clara, CA). Their concentrations were determined using the QX200 digital PCR system (Bio-Rad, Hercules, CA). Libraries were pooled in the same sequencing run at specific ratios: 95% mRNA, 5% ADT. Sequencing was performed on the NextSeq2000 (Illumina, San Diego, CA) using the 50-bp P3 NextSeq kit. The cycle distribution was: [read 1 (28 cycles), i7 index (8 cycles), and read 2 (52 cycles)]. Sequencing depth exceeded 50,000 reads/cell.

### Antibodies used for scRNA-seq

We used a range of TotalSeq (BioLegend) oligo-tagged antibodies to complement our mRNA-targeted scRNA-seq analysis. The clones for these antibodies were validated for cross-specificity against *M. fascicularis* commercially by BioLegend, in the published literature, or in-house by researchers using NHPs at the University of Pittsburgh. These antibodies were added to our ADT antibody cocktail mix, each at a concentration of 0.5 ug/sample, and included antibodies against CD1c, CD1d, CD3, CD4, CD8, CD11b, CD11c, CD14, CD16, CD36, CD63, CD64, CD80, CD86, CD103, CD163, CD195, CD206, CD273, CD274, CD279, and HLA-DR. Details including the clone, host, and product number for each antibody are included in Supplementary Table 2.

### Immunofluorescence microscopy for visualizing Mtb localization and macrophage subsets

Bacteria and cell subsets were visualized in formalin-fixed paraffin-embedded tissue (FFPE) sections. FFPE blocks were prepared by the University of Pittsburgh Division of Laboratory Animal Resources (DLAR) and 5 um-thick tissue sections were cut by DLAR histology technicians. For immunofluorescence-based immunohistochemistry (IHC), the slides were deparaffinized with sequential dips in xylenes (2x) and 95% EtOH (2x), then washed with deionized water (dH_2_O) to remove any residual alcohol. A cyclic IHC process similar to that previously published[11, 31] was used to enable multiple rounds of staining on the same tissue section. Briefly, after the slides were deparaffinized, antigen retrieval was performed in Tris-EDTA-Tween buffer (pH 9.0) as previously indicated[31, 32] and the slides were blocked with 1% BSA-PBS for 30 minutes at RT. Primary antibody cocktails were designed to include multiple mouse isotypes and a rabbit antibody so that isotype-specific mouse secondary antibodies could be used in the same secondary cocktails, thus enabling multiple labeling with antibodies from the same host species. The antibodies and related products used in this study are listed in Supplementary Table 4. Slides were incubated with primary antibodies for 60 minutes at RT or overnight at 4°C, and then primary antibodies were labeled with fluorochrome-conjugated secondaries. Zenon labeling (Thermo Fisher Scientific, Waltham, MA) was used as a tertiary step to expand panels where multiple mouse antibodies with the same isotype were included in a panel. After the final round of staining, the slides were washed with PBS and coverslips were applied with DAPI-containing Prolong Gold Mounting Medium (Thermo Fisher Scientific).

Stained tissue sections were visualized with a Nikon e1000 epifluorescence microscope (Nikon, Melville, NY) with a 20x objective (Nikon Apo 0.75 NA) using a DS-Qi2 digital camera (Nikon) operated by Nikon Elements AR imaging software version 4.5. All the images were acquired as stitched images with 10% overlap between imaging fields that were processed with Nikon Elements AR software. Images were saved as 14-bit ND2-format files. After imaging, the coverslips were removed by incubating the slide in deionized H2O in a Coplan jar until the coverslip came off and the slides were washed with PBS to remove residual mounting medium. The previous round of antibodies were stripped by subjecting the slides to a second round of antigen retrieval followed by stripping with the Vecta Plex Antibody Removal Kit (Vector Laboratories, Newark, CA) per the manufacturer’s instruction. Efficient stripping was visually confirmed before the slides continued in the staining process. After the stripping was complete, slides were blocked with 1% BSA-PBS for 30 minutes and processed as previously indicated. After the secondary or tertiary staining was complete, autofluorescence was quenched using the Vector TrueVIEW Autofluorescence Quenching Kit (Vector Laboratories) per manufacturer protocols and the coverslips were applied using DAPI-containing mounting medium and imaged as previously noted. Slides were stored at -20°C between rounds of staining. QuPath [33] was used to segment cells and to capture their x-y coordinates in the images so the position of marker-positive cells could be plotted in the granuloma’s spatial organization with CytoMAP [34]. Adobe Photoshop version 2024 (Adobe, San Jose, CA) was used to prepare figures using TIFF-format files exported from the original ND2 files and for combining the IHC images with the mapped cells.

## Data Analysis

### Data Acquisition and QC

Sequencing data derived from each run were processed using software tailored to the library type. mRNA libraries were processed using CellRanger software (v. 6.0) (10X Genomics), utilizing a custom reference genome combining the *M. fascicularis* 5.0 assembly (GCF_000364345.1) and the Mtb Erdman genome (GCA_00668235.1). ADT libraries were processed using CITE-Seq-Count (v. 1.4.3), available at https://hoohm.github.io/CITE-seq-Count/. This processing yielded raw count matrices for both mRNA and proteins. Downstream data analysis was conducted in Seurat (v. 4.3) [35]. To ensure robust data, cells with fewer than 700 unique genes as well as cells with a high proportion (>5%) of ribosomal protein genes (RPS and RPL) were identified and removed.

### Single-Cell RNA Sequencing Data Preprocessing and Integration

In our scRNA-seq analysis, we processed individual samples’ datasets using Seurat’s regularized negative binomial regression, regressing out the number of counts, as in Hafemeister and Satija [36]. Metadata columns such as ‘*Batch*,’ ‘*Type*,’ and ‘*Infection’* were then added to each dataset. Next, we combined all datasets into a single Seurat object. The raw counts from the RNA slot of this merged object served as input for Harmony integration [37]. Raw counts underwent log-normalization, followed by the identification of the top 5,000 variable genes. These genes were then scaled and centered. Subsequently, we performed Principal Component Analysis (PCA) on these values. Finally, we performed integration with Harmony, using ‘Batch,’ ‘Infection,’ and ‘Type’ as covariates (Supp. Figure 3A).

### Cluster Detection and Annotation

Using the Harmony-aligned embeddings, graph-based cluster detection was achieved utilizing principal components (PCs) that had been marked statistically significant by the jackstraw method [38]. For community detection, we employed the Louvain algorithm [39]. Both reference-based approaches [40] and canonical markers gene analysis were used to annotate the cell type of each identified cluster.

### Trajectory and Pseudotime Analysis

This analysis was carried out in Monocle (v3.0)[41] using the integrated Seurat object, which was converted with the SeuratWrappers package in R (v. 0.2.0). For unbiased trajectory and pseudotime analysis of the macrophage populations, all cells classified as macrophages were assigned to the same partition, and trajectory/pseudotime analysis was conducted as previously described[42]. White circles indicate the root origins of the trajectory, grey circles indicate destination fates and black circles indicate branching points.

To investigate genes that vary over pseudotime around branching point 1, cells around the branching point were selected using the *choose_cells()* function. The subsetted dataset was preprocessed with the *preprocess_cds()* function, which normalizes the data, calculates a lower-dimensional space, and prepares the data for trajectory inference. Significant genes were identified using the *graph_test()* function, which tests for differential expression based on the low-dimensional embedding and the principal graph. Gene modules with significant variation in expression around branching point 1 were identified using the *find_gene_modules()* function and aggregated using the *aggregate_gene_expression()* function.

### Weighted Gene Co-expression Network Analysis (WGCNA)

To perform weighted gene co-expression network analysis on our scRNA-seq dataset, we employed the scWGCNA package [43]. Initially, 20,000 cells were randomly subsetted from the original dataset without replacement. Pseudocells were then computed using the *calculate.pseudocells* function from the scWGCNA package. This method aggregates single cells into pseudocells to reduce the complexity of the dataset. A fraction of 0.2 of the cells were used as seeds, and for each seed, 10 nearest neighbors were aggregated based on the Harmony dimensional reduction. The single cell data were normalized using the *LogNormalize* method with a scaling factor of 10,000. Variable features were identified using the Variance Stabilizing Transformation (VST) method, and the top 3,000 features were retained.

Subsequently, pseudocells and the variable genes identified in the previous step were used in the *run.scWGCNA* function to identify modules of co-expressed genes. Membership tables were inspected to understand the genes belonging to each module, and average expressions of each co-expression module per cell were analyzed. Eigengenes for the co-expression modules were computed using the *scW.eigen* function, which collapses the expression profile of each module into a single representative profile known as the module eigengene. The average expression of each module was then visualized on a UMAP plot using a customized *scW.p.expression* function for improved visualization.

### Differential Abundance Analysis using MiloR

Compositional analysis of cell populations between granuloma and non-diseased lung was conducted using the MiloR package [44]. A k-nearest neighbors (k-NN) graph was constructed using the *buildGraph* function with *k*=10, employing Harmony for dimensionality reduction. Neighborhoods were defined on the graph using the *makeNhoods* function, with the same *k* and *d* parameters. To optimize the model parameters, both the neighborhood cell distribution and the distribution of uncorrected P-values were assessed. Cells within these neighborhoods were counted based on their originating samples using the *countCells* function. Differential abundance testing of the neighborhoods was performed using the *testNhoods* function, incorporating a design matrix that included sample identifiers, batch information, and a tissue type variable. The results were sorted by the spatial False Discovery Rate (SpatialFDR).

A neighborhood graph was further constructed using the *buildNhoodGraph* function. Visualizations of the neighborhood graph, highlighting the differential abundance results, were generated (Figure 5A). Neighborhoods were then annotated based on cell identity, with those having an identity fraction less than 0.8 labeled as “Mixed”. Customized bee swarm plots were created to visualize the data based on these identities (Figure 5B). Finally, neighborhoods were grouped using the *groupNhoods* function with a *max.lfc.delta* of 2. The grouped neighborhoods were visualized on a UMAP plot and further explored through customized bee swarm plots based on their LFC differences (Figure 5C and Supp. Figure 6A).

### Gene Set Enrichment Analysis (GSEA)

Gene Set Enrichment Analysis (GSEA) was performed to compare the expression profiles of AMs and IMs. All AM cells were combined into a single cluster, and their expression profile was compared against the combined population of all IM cells. Differential expression analysis was conducted to generate a ranked list of genes based on the formula *SIGN(LogFC)*-LOG10(p-value)*, where LogFC represents the log fold change, and p-value represents the significance of the differential expression.

The ranked gene list was imported into GSEA version 4.2.3. The GSEApreranked test was conducted using the hallmark gene sets (H:) from the human molecular signature database (MSigDB). The parameters for the analysis were set as follows: 2,000 permutations, enrichment statistic set to weighted, excluding gene sets larger than 500 genes and smaller than 10 genes. This analysis allowed the identification of significantly enriched gene sets in the context of the differential expression between AM and IM populations.

### Protein-Protein Interaction Network Analysis

A protein-protein interaction (PPI) network analysis was conducted using Cytoscape to explore the interactions among genes associated with the TR-AM population. A total of 869 genes from module 11, identified through the scWGCNA analysis, were imported into Cytoscape using the STRING app. A confidence score cutoff of 0.7 was applied to generate the PPI network.

### Statistical Analysis scRNA-seq

Differential expression analysis was conducted using the nonparametric Wilcoxon rank-sum test as implemented in Seurat [45]. Only genes with a false discovery rate (FDR) of less than 0.05 between two comparisons were considered statistically significant. Unless specified otherwise, the Wilcoxon rank-sum test followed by FDR correction was used to compare gene expression distributions between two groups of cells in various plots and visualizations.

In addition, effect sizes for differences in expression between groups were quantified using Cliff’s Delta, which measures the probability that a randomly selected value from one group will be greater than a randomly selected value from the other group. Cliff’s Delta values range from -1 to 1, where 0 indicates no effect, and values closer to -1 or 1 indicate stronger effects. The Cliff’s Delta was computed using the *cliff.delta()* function from the effsize R package (available at doi:10.5281/zenodo.1480624).

Data integration and batch effect correction were performed using Harmony, following the method previously described[37]. ADT and HTO data were normalized using a centered log ratio transformation via the *NormalizeData()* function with *normalization.method = ‘CLR’* in Seurat. RNA counts were log-normalized and scaled prior to principal component analysis (PCA) and data integration with Harmony. Data visualizations, including feature plots, scatter plots, and violin plots, were generated using log-normalized counts. Heatmaps and dot plots were created using scaled expression data, as per the default settings in Seurat.

## Results

### Characterization of alveolar and interstitial macrophages in the lung and granulomas of Mtb-infected macaques

We aimed to characterize the phenotypic and functional diversity of macrophages in TB granulomas using a multi-modal approach combining scRNA-seq, immunofluorescence staining, and flow cytometry (Figure 1A). To achieve this, we infected three Mauritian cynomolgus macaques with an mCherry-expressing *M. tuberculosis* (Mtb) strain, to track infected cells and facilitate their sorting via flow cytometry, as previously described[13, 17] (Supplementary Figure 1). Lung infection was confirmed at four weeks post-infection by PET-CT imaging, which revealed multiple granulomas in each animal (Figure 1B); note that this is a very early time point in Mtb infection when granulomas are newly formed. These granulomas displayed a range of morphologies, including both caseous (necrotic) and non-caseous structures (Figure 1C).

**Figure 1.**
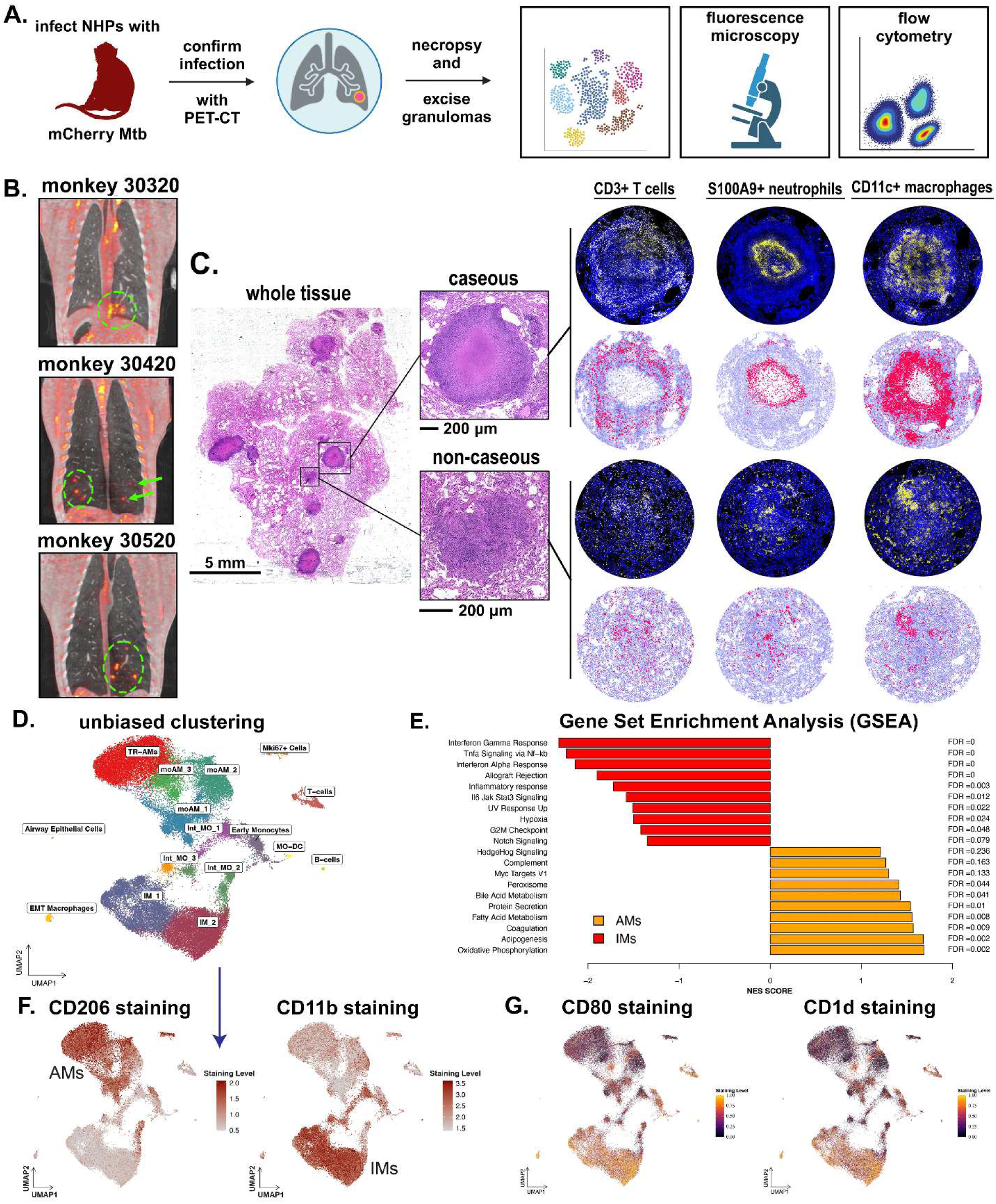
NHP infection with mCherry-expressing Mtb leads to TB disease where macrophages can be interrogated by scRNAseq. A. Schematic of the overall experimental design. NHPs were infected with mCherry-expressing Erdman-strain Mtb and the infection was confirmed using PET-CT scans. At necropsy, granulomas and non-diseased lung tissues were excised and frozen for multimodal scRNA sequencing, fluorescence microscopy, and flow cytometry for comparative analysis. B. ^18^F-FDG PET-CT imaging shows the development of TB-associated inflammation in the lower lung lobes of monkeys 30320, 30420, and 30520. Arrows and circles indicate the locations of granulomas. C. Diseased lung tissue from monkey 30520 shows numerous lesions, including caseous and non-caseous granulomas, with positive staining for CD3^+^ T cells, S100A9^+^ neutrophils, and CD11c^+^ macrophages (yellow) and DAPI-stained nuclei (blue). Immunofluorescence images for each marker (left) were used to identify the location of marker-positive cells (red) against a background of marker negative cells (gray). D. UMAP visualization of scRNA-seq data from 51,214 cells collected from both granuloma and non-diseased lung tissue. Cells are clustered in an unbiased manner based on their transcriptional profiles. E. GSEA of differentially expressed genes between AM and IM, using the hallmark gene sets (H:) from the Molecular Signatures Database of the Broad Institute. The figure displays the normalized enrichment scores (NESs) and false discovery rates (FDRs) for each gene set, highlighting the pathways that are most significantly enriched in AMs and IMs. F. Immunostaining of lung cells shows differential expression of CD206 and CD11b between AMs and IMs, corresponding to the ontologically and transcriptionally defined characterization of these macrophages. G. UMAP visualizations of cells stained for CD80 and CD1d, with normalized staining levels represented by a color gradient from black (low) to yellow (high).

We hypothesized that macrophage populations would differ between non-diseased (healthy) and diseased (granulomatous) lung tissues. To test this hypothesis, we cryopreserved samples from both healthy lung areas and TB granulomas (Supp. Figure 2 – Panels A-D), to be analyzed separately and in an integrated manner. Following cryopreservation, tissues were thawed, mechanically disaggregated, flow- and ADT-stained using a panel of 22 oligo-tagged antibodies (Supplementary Table 4) to label surface markers [30]. We then used flow cytometry to sort macrophages based on forward and side scatter enrichment and mCherry fluorescence, separating infected (mCherry^+^) and uninfected (mCherry^-^) populations (Supplementary Figure 1). The sorted cells were subsequently prepared for scRNA-seq library construction.

After sequencing, data integration, and quality control (Supp. Figure 3A), we recovered 51,214 cells from three animals for downstream analysis. Unbiased clustering of the transcriptomic data identified 16 distinct subpopulations, most of which were macrophages, as expected. In addition to macrophages, we detected smaller populations of T cells, B cells, and epithelial cells in the lung samples (Figure 1D). For the purposes of this study, we focused on the macrophage clusters to define their phenotypic and functional diversity and to understand their roles in the early immune response against Mtb.

Reference-based analysis[46] of the scRNA-seq data distinctly delineated the two main ontological macrophage populations in the lung: alveolar (AMs) and interstitial macrophages (IMs) (Supp. Figure 3B). To explore the functional properties of AM and IM in NHPs lungs, we performed Gene Set Enrichment Analysis (GSEA) on their differential transcriptomes (Supplementary Data 1). Consistent with previous studies[47], GSEA revealed that AMs exhibited metabolic polarization towards fatty acid metabolism and oxidative phosphorylation, pathways associated with tissue homeostasis but also implicated in facilitating Mtb persistence and replication[17]. In contrast, IMs were characterized by expression of pro-inflammatory pathways, including interferon-gamma (IFN-γ), tumor necrosis factor (TNF), and NF-κB signaling (Figure 1E). Oligonucleotide-tagged immunostaining identified CD206 and CD11b as key surface markers that distinguish AMs from IMs, respectively, (Figure 1F) and flow cytometry analysis confirmed the presence of both AM and IM subsets in the lung samples (Supplementary Figure 3C). Complementing these findings, we observed differential staining for co-stimulatory and antigen-presenting molecules between the two macrophage subsets. Notably, CD80 – a co-stimulatory protein required for T-cell activation via CD28 and a marker of classically activated (M1) macrophages previously associated with drug tolerance induction in Mtb[48], and CD1d, a lipid antigen-presenting molecule to CD1d-restricted T-cells such as NKT cells[49], were expressed by the pro-inflammatory IMs but not AMs (Figure 1G).

### The macrophage landscape in early Mtb infection is dominated by extravasating monocytes undergoing polarization

The heterogeneity observed across AM and IM populations, as revealed by UMAP analysis (Figure 1D), prompted us to investigate their ontogeny and differentiation pathways. Previous studies suggest that human CD14^+^ and CD16^+^ monocytes extravasate into the lung and differentiate into diverse macrophage populations, including both AMs and IMs[50]. Using CITE-SEQ surface staining, we leveraged CD14 and CD16, markers of monocyte-derived cells, to show that most macrophages in infected NHP lung tissue were monocyte-derived, with the exception of tissue-resident alveolar macrophages (TR-AMs) (Figure 2A – Supp. Figure 4A). This monocyte origin was further corroborated by the expression of *Mef2c*, a gene associated with monocytic differentiation[51], and *Csf1r*, a marker of bone marrow-derived myeloid cells[52–55] (Figure 2B). To further investigate the transcriptional programs driving the origins and differentiation of monocyte-derived macrophages during early infection, we performed single-cell weighted gene co-expression network analysis (scWGCNA) on the 51,214 cells in our dataset[43]. This analysis identified 23 distinct gene co-expression modules (Supplementary Figure 4B), many of which recapitulate the origin and function of specific macrophage subsets. In particular, one module consisting of 25 co-expressed genes was expressed in both monocyte-derived dendritic cells (MO-DCs) and a subset of cells within the “Early Monocytes” cluster, which we termed “Blood and Recruited Extravasating Monocytes” (Figure 2C, Supplementary Data 2). This module contained DC markers such as *Xcr1* and *Clec9a* (Figure 2D, Supp. Figure 4C)[56–59], along with genes involved in cell-cell adhesion and leukocyte migration, including CDH20 (Figure 2E), PAK3, TSPAN33, and CADM1[60–63]. The “Early Monocytes” cluster also expressed CPVL, a gene involved with lamellipodium formation, alongside additional regulators of cell migration and chemotaxis—DOCK10, PAK1, RGS1, RGS12, and ALCAM (Figure 2F). Collectively, these data suggest that these cells originate from common monocyte-DC progenitors in peripheral blood, have recently extravasated into infected lung tissue, and are undergoing differentiation within the granuloma microenvironment[64–68].

**Figure 2.**
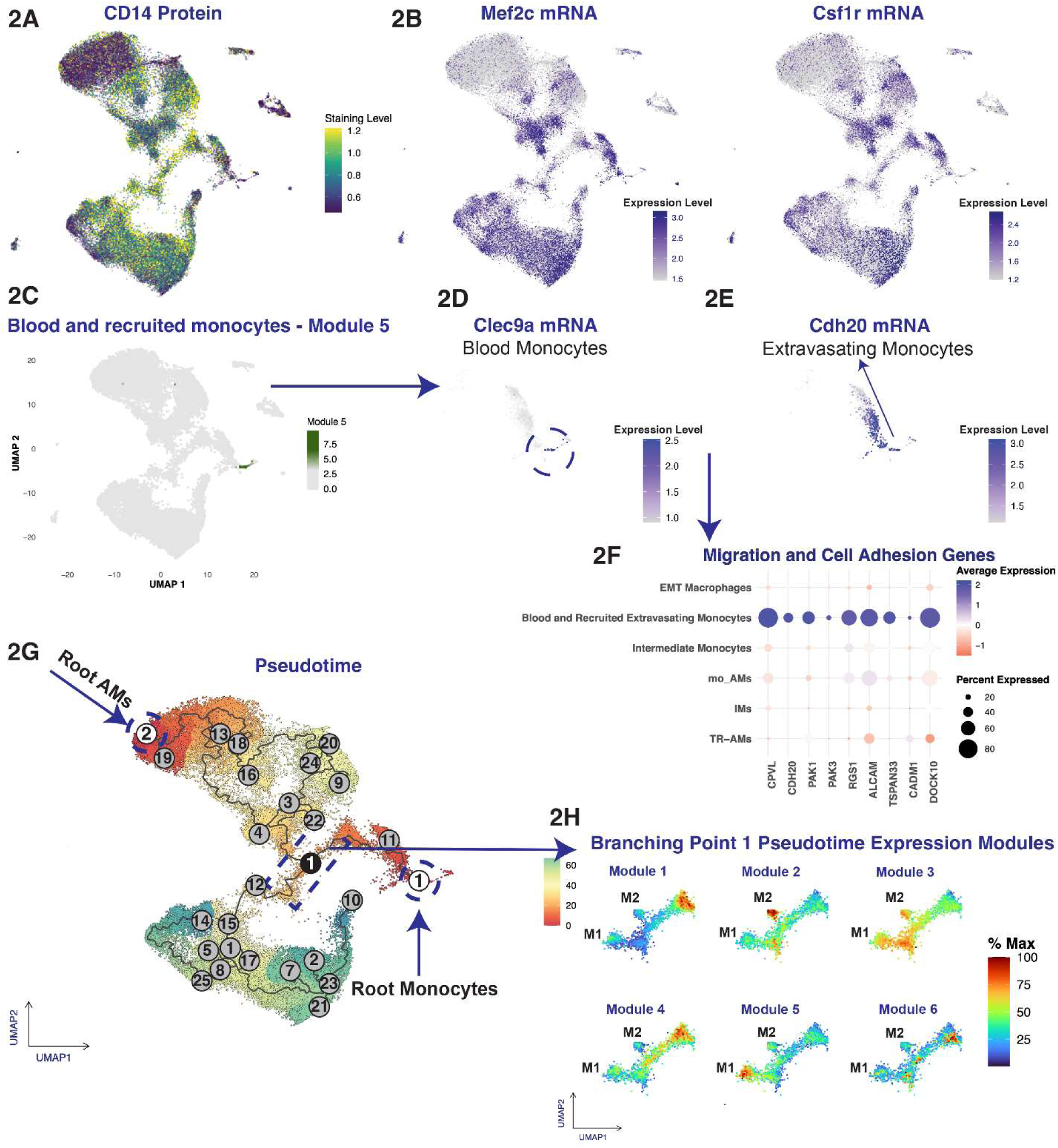
Monocyte extravasation and differentiation in the Mtb-infected lung. A. UMAP plot displaying CD14 antibody staining intensity across single cells, highlighting the lineage-specific staining pattern of CD14 within monocyte populations. B. UMAP plots showing log-normalized expression levels for the *Mef2c* (left) and *Csf1r* (right) genes across all cells. C. Visualization of scWGCNA-derived Module 5 (25 genes) expression levels mapped onto UMAP embeddings of all cells. This gene module is primarily expressed in blood and extravasating monocyte subpopulations. D. Differential expression of key markers, *Clec9a* and *Cdh20*, associated with blood presence and extravasation processes within the Early Monocyte subset. The UMAP plots spatially represent the early phases of monocyte differentiation, from their arrival into the lung to their extravasation into the tissue. E. DotPlot showing expression levels of key migration and cell adhesion genes across various macrophage populations. Dot size represents the percentage of cells expressing each gene, while color intensity indicates the average expression level. F. Trajectory and pseudotime analysis of macrophages visualized on the UMAP projection. The inferred differentiation trajectory is shown, with starting points (“roots,” white), terminal states (“leaves,” gray), and intermediate branching points (black) where the trajectory bifurcates. The numbers label each key point (roots, leaves, and branching points) and correspond to identifiers assigned by Monocle during trajectory reconstruction. G. Identification and analysis of gene expression modules around Branching Point 1, a critical juncture in the monocyte-to-macrophage differentiation process. The analysis focuses on cells clustered near this branching point, identifying gene modules with significant expression changes across the pseudotime axis. UMAP visualization displays these gene modules and provides insights into their spatial distribution at this transitional stage, with a color scale indicating the intensity of expression.

Trajectory and pseudotime analyses further supported these findings by revealing the differentiation paths of recruited monocytes. The analysis identified a trajectory rooted in XCR1^+^ and CLEC9A^+^ monocytes, which branches at the key bifurcation point 1 (denoted by a black circle in Figure 2G). From this juncture, monocytes follow one of two distinct routes: they either differentiate toward M2-polarized AM phenotypes (indicated by gray circles) or progress toward M1-like IMs (Figure 2G). This bifurcation suggests that distinct microenvironmental cues within infected lung tissue guide monocytes into either an anti-inflammatory (M2) or a pro-inflammatory (M1-like) state during early Mtb infection.

To investigate the signals driving differentiation around branching point 1, we targeted cells near this bifurcation and analyzed their cellular dynamics and gene expression changes along the pseudotime axis. Six gene modules with unique expression profiles emerged near the split (Figure 2H; Supplementary Data 3). Module 1 is associated with early-stage monocytes, enriched for pathways regulating protein synthesis, RNA processing, and other homeostatic functions (Supplementary Figure 4D). Module 4, expressed by cells transitioning from early monocytes to branching point 1, is linked to immune activation pathways—cytokine signaling, Toll-like receptor cascades, TNFR1 signaling, and MyD88-independent TLR4 pathways—critical for early immune responses following infection (Supplementary Figure 4E). Module 3 extends from branching point 1 toward M1 macrophages (though some expression overlaps with M2 cells) and includes pathways connected to membrane trafficking, vesicle-mediated transport, receptor tyrosine kinase signaling (e.g., VEGF), and RHO GTPase activity, which mediate cell motility, survival, and proliferation (Supplementary Figure 4F). In contrast, Module 2 is associated with monocyte differentiation into monocyte-derived alveolar macrophages (moAMs), enriched for pathways linked to cytoskeletal dynamics and cell migration (e.g., Rho GTPase, RAC1). Additionally, the expression of RORA and genes involved in mitochondrial biogenesis suggests a transition toward tissue-resident, M2-like anti-inflammatory states (Supplementary Figure 4G)[69, 70]. Lastly, Module 5, expressed in cells that differentiate toward M1 macrophages, is tied to pro-inflammatory responses, cell survival, and apoptosis, featuring TNF, MAPK, and FoxO signaling pathways that regulate inflammatory gene expression and stress responses (Supplementary Figure 4H)[71–74].

Overall, these findings reveal that the early phase of pulmonary TB is marked by extensive remodeling of the myeloid cell landscape, driven by peripheral blood monocytes that differentiate locally within the lung to form polarized macrophage subsets. These subsets collectively represent the predominant macrophage populations during this stage of infection, underscoring their importance in shaping granuloma development and pathogen control.

### Immunostaining reveals the heterogeneity of AM subpopulations in Mtb-infected lungs

Focusing on the AM lineage, unbiased UMAP clustering identified four distinct subpopulations, underscoring the heterogeneity of AMs in Mtb-infected lung tissue (Figure 1D). Immunostaining revealed differential expression of canonical and alternative surface markers that distinguish monocyte-derived alveolar macrophage subsets (moAMs_1-moAM3) from tissue-resident alveolar macrophages (TR-AMs). Expression of CD206, a hallmark AM receptor, varied notably in moAMs depending on whether they originated from granuloma-containing or non-diseased lung tissue. This variability was mirrored by changes in CD63 levels, suggesting that moAMs undergo phenotypic reprogramming in response to localized inflammatory conditions within granulomas (Figure 3A, Supp. Figure 5B). Additional distinctions emerged from CD86 staining—a CD28 ligand indicative of antigen-presenting capabilities and immune activation[75]—which was predominantly expressed by moAMs, distinguishing them from TR-AMs (Figure 3B). In contrast, CD163, a marker of alternatively activated (M2-like) macrophages, labeled TR-AMs in non-granulomatous lung tissue (Figure 3C, 1D), but was detected in both moAMs and TR-AMs in granulomas (Supp. Figure 5A). Meanwhile, CD195 (CCR5)—a chemokine receptor involved in inflammatory responses[76, 77]—and CD14, an LPS receptor[78], were observed exclusively in monocyte-derived cells, highlighting the interplay of ontogeny and polarization in defining macrophage phenotypes[12] (Figures 3B, 2A).

**Figure 3.**
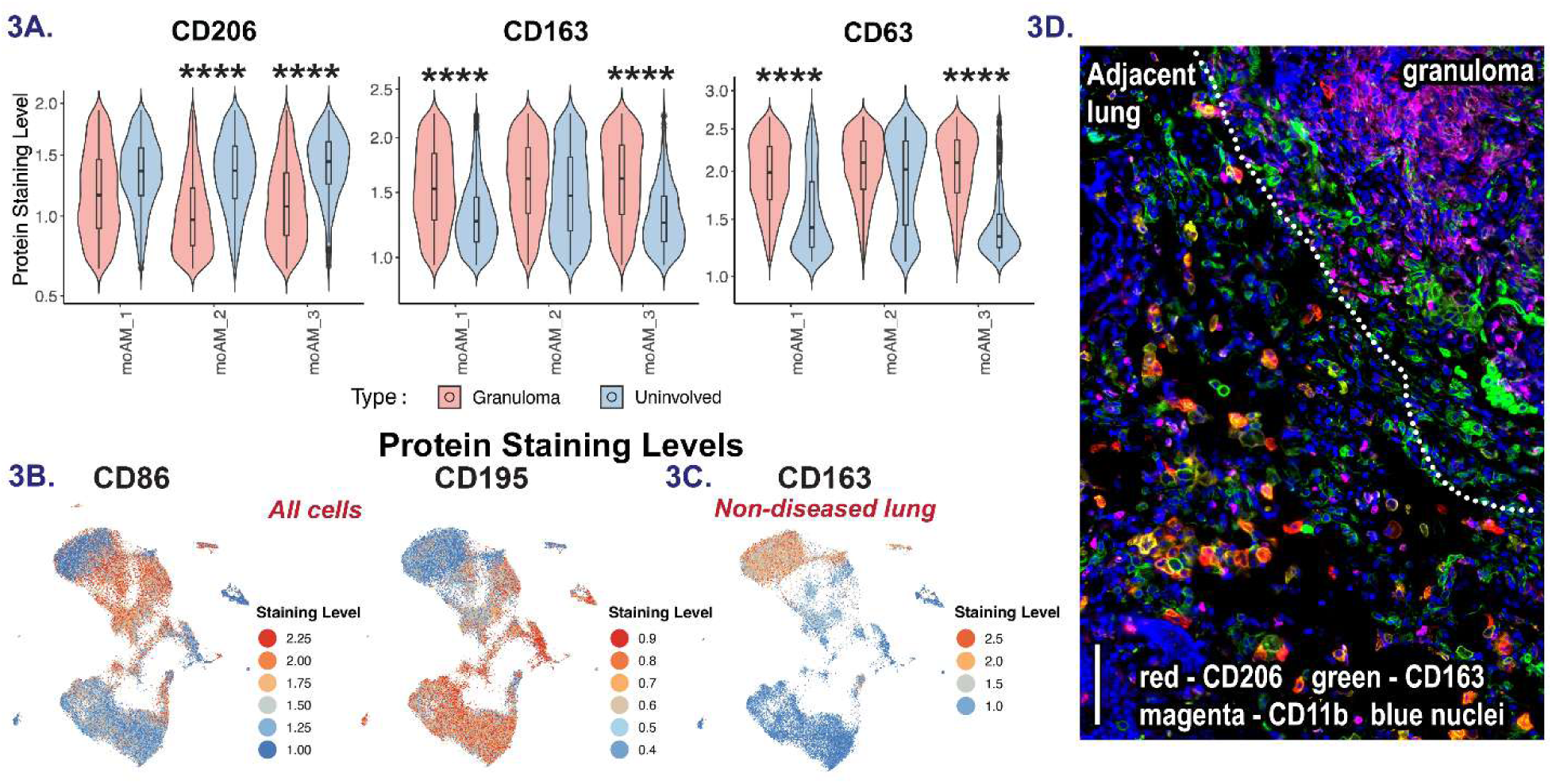
MoAM subsets differ between granuloma and non-diseased lung. A. CD206, CD163, and CD63 surface marker staining of moAM subsets within granuloma and non-diseased lung with moAM numbering (moAM_1, moAM_2, moAM_3) corresponding to the moAMs identified through UMAP-based clustering analysis in Figure 1D. Data are presented as violin plots overlaid with box plots. The center line of each box plot represents the median expression level of the marker, the box bounds represent the interquartile range (IQR) from the 25th to 75th percentile, and the whiskers extend to 1.5 times the IQR. Statistical significance was assessed using a two-sided Wilcoxon Rank-Sum Test to compare expression levels between ‘Granuloma’ and ‘Uninvolved’ cell types within each moAM subset. P-values were adjusted for multiple comparisons using the Benjamini-Hochberg procedure, with significance indicated by markers for extremely low adjusted p-values (**** = p.adj < 1e^-50^). B. UMAP visualizations of cells stained for CD86 and CD195 protein markers. Each plot uses a color gradient from dark blue (low expression) to dark red (high expression) to represent staining intensity across a spectrum of expression percentiles. C. UMAP plot illustrating staining levels for the CD163 marker, a hemoglobin scavenger receptor, across non-diseased lung tissue. The plot uses a color gradient from dark blue (low expression) to dark red (high expression) to represent staining intensity across a spectrum of expression percentiles. D. Representative visualization of IHC staining for surface markers CD206, CD163, and CD11b at the granuloma edge. The staining patterns reveal distinct spatial distributions: CD206^+^ CD163^-^ and CD206^+^ CD163^+^ moAMs and TR-AMs predominate in areas adjacent to the granuloma, while CD163^+^ CD206^-^ moAMs are primarily located in the inner core. Additionally, CD11b^+^ IMs are present within the core. Scale bar (left side bottom, vertical) = 100 μm.

To further explore the complexity of AM populations in relation to granuloma proximity, we used immunohistochemistry (IHC) to visualize CD206/CD163-expressing AM subsets alongside CD11b-expressing IMs in granulomas (Figure 3D). We found that granuloma-adjacent lung tissue contained a high prevalence of CD206^+^ AMs, while AMs closer to granulomas primarily expressed CD163 with reduced or absent CD206, indicating an activated moAMs profile (see later)(Supplementary Figures 5A, 5B). Importantly, CD163^+^ monocyte-derived cells with decreased CD206 expression are prevalent in lungs from macaques with active TB compared to those with latent TB exhibit IFN-responsive phenotypes, and are identified by their expression of ligands for the PD-1 receptor and CD38 [7]. Our scRNA-seq data, combining RNA expression and surface protein detection via CITE-seq, showed that moAMs expressed mRNA and protein for the PD-1 ligand CD273 (PD-L2) in granulomas and have increased CD38 RNA expression — a recently defined marker of pro-inflammatory macrophages in murine TB[12] (Supp. Figures 5C-5D). Both CD206^+^ and CD163^+^ AMs were absent from the granuloma’s IM-rich myeloid core (Figure 3D), suggesting that their main influence on granuloma dynamics is localized to the peri-granuloma region and outer lymphocyte cuff. Overall, these findings emphasize the plasticity and complexity of the AM population in Mtb-infected lung tissue, indicating moAMs undergo inflammatory reprogramming in granulomas and potentially suggesting distinct roles for TR-AM and moAMs in shaping granuloma structure and function.

### Characterization of moAMs and TR-AMs in non-diseased lung tissue

Building on our phenotypic characterization of AM subsets, we next investigated the transcriptional programs that define the ontogenic origins of moAMs and TR-AMs. To exclude transcriptional signals driven by granulomatous inflammation, we performed scWGCNA analysis on 24,735 cells isolated from non-diseased lung tissue. This analysis identified two gene co-expression modules matching the protein-level staining profiles of moAMs and TR-AMs. Specifically, CD206^+^CD163^-^ moAMs were defined by a 138-gene module (module 3) (Figure 4A), which contained monocyte-specific markers and genes linked to monocyte-to-macrophage differentiation (Supplementary Figure 5E, Supplementary Data 4). Notably, this subset expressed the transcription factor Maf, a key regulator of M2-like homeostatic and reparative macrophage phenotypes[79–81] (Figure 4B). The elevated expression of Maf-regulated genes suggests a homeostatic role for these moAMs in non-granulomatous lung tissue (Figure 4C), and the ability of IFN-γ, highly present in the context of granulomas (see later), to repress M2 gene expression by disassembling enhancers bound by Maf[80], likely explains their ability to undergo pro-inflammatory reprogramming in these lesions.

**Figure 4.**
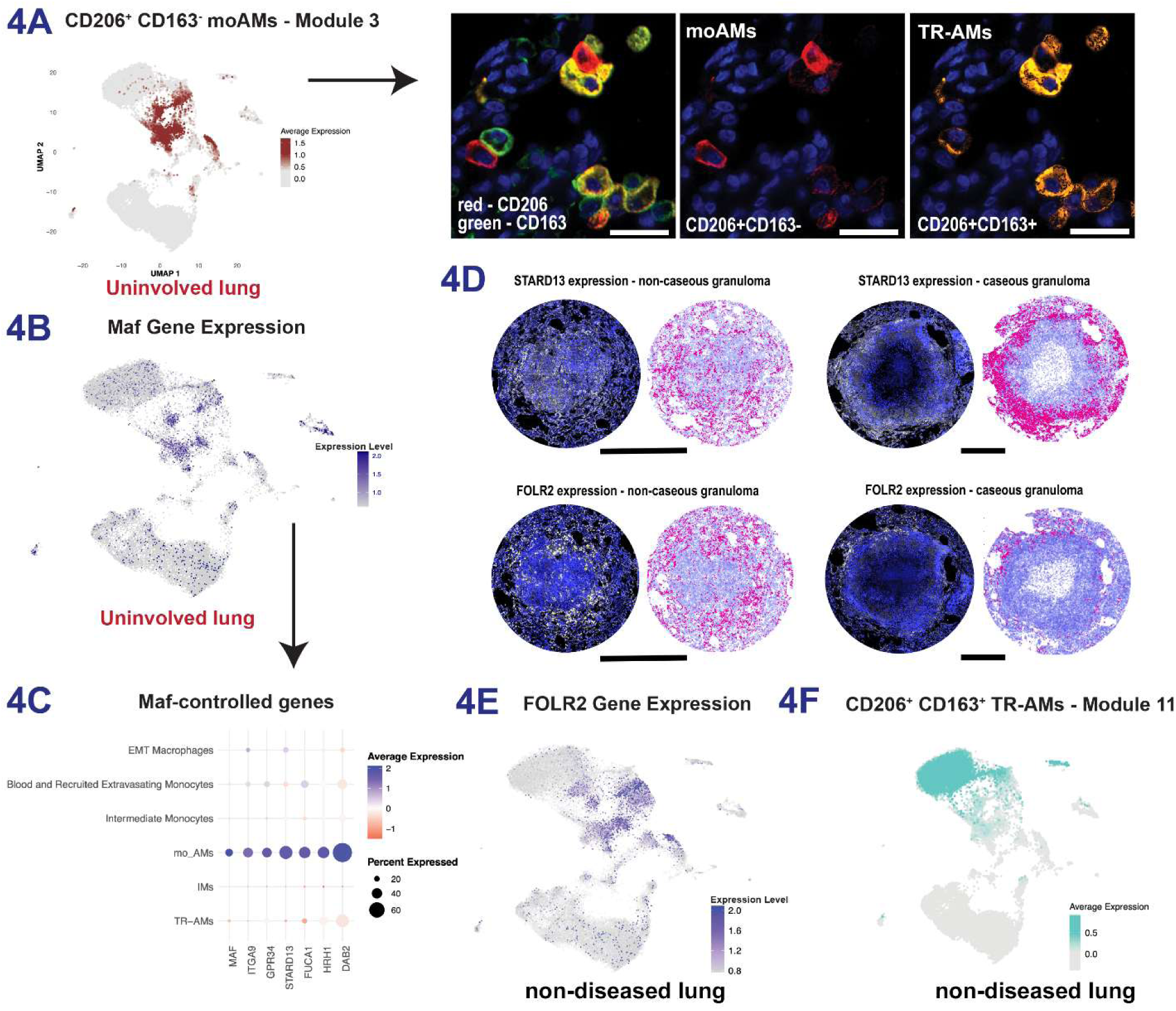
MoAMs are transcriptionally and phenotypically distinct from TR-AMs. A. Visualization of average expression levels of the scWGCNA-derived Module 3 (139 genes), specific to CD206^+^ CD163^-^ moAMs, mapped onto UMAP embeddings from non-diseased cells. Scale bar = 50 μm. B. UMAP plot showing log-normalized expression levels of the *Maf* gene among non-diseased lung cells. C. DotPlot displaying expression levels of *Maf*-controlled genes across various macrophage clusters. Dot size represents the percentage of cells expressing each gene, while color intensity indicates the average expression level. D. Localization of cells expressing STARD13 and FOLR2 in a non-caseous (left) and caseous (right) granuloma. Each paired image includes the microscopy-based image (left half) and the plotted positions of the positive cells (red) against the total cell population (blue) (right half). Scale bar = 500 μm. E. UMAP plot showing log-normalized expression levels of the *Folr2* gene across all cells. F. Visualization of average expression levels of the scWGCNA-derived Module 11 (869 genes), specific to CD206^+^ CD163^+^ TR-AMs, mapped onto UMAP embeddings from non-diseased cells.

To determine the spatial localization of these cells, we visualized STARD13, a gene in this module, within both non-caseous and caseous granulomas, finding STARD13^+^ cells predominantly at the granuloma periphery (Figure 4D, top). Given that the granulomas included some cellular structures from the lung tissue that was immediately adjacent to the granulomas, the STARD13⁺ cells in the outer areas likely reflect macrophages from the surrounding lung tissue. Similarly, we observed FOLR2 expression in moAMs (Figure 4E), a marker previously linked to tissue-resident and M2/anti-inflammatory macrophages[82, 83]. Like STARD13^+^ cells, FOLR2^+^ moAMs localized to the lung tissue adjacent to granulomas and the lymphocyte cuff (Figure 4D, bottom). As previously noted, their monocyte origin was confirmed by CD14 staining and expression of Mef2c and Csf1r (Figures 2A, 2B).

In contrast, CD206^+^CD163^+^ TR-AMs—which are embryonically derived—were defined by an 869-gene module (Figure 4F). This module included canonical TR-AM markers (CD36, FABP4, LPL, and SCD[47]) (Supplementary Figure 5F; Supplementary Data 5). Pathway and protein–protein interaction analyses of this gene set revealed enrichment for fatty acid metabolism, oxidative phosphorylation, TGF-β signaling, and cell adhesion, aligning with previous studies (Supplementary Figure 5G)[17, 47]. Altogether, these findings delineate two principal AM populations with homeostatic functions, both localized near granulomas. During early pulmonary TB, these populations can undergo reprogramming in inflammatory environments, altering their surface marker expression and potentially contributing to immune regulation and tissue remodeling.

### Increased abundance of AMs with inflammatory phenotypes in granuloma

To characterize the extent of AM reprogramming in inflammatory conditions, we next examined macrophage heterogeneity in the context of overt disease by evaluating how granulomatous inflammation influences local macrophage populations. Using differential abundance testing[44] (Figure 5A), we compared the composition of macrophages in granulomatous versus non-diseased lung tissue (Figure 5B). A neighborhood-based grouping strategy revealed subpopulations either enriched in granulomas (groups 6, 2, 8) or underrepresented (group 3) (Figure 5C). Group 6, which showed the largest increase in granulomas, predominantly comprised multiple moAM populations, including moAM_2, moAM_3, alongside a subset of TR-AMs (Supplementary Figure 6A).

**Figure 5.**
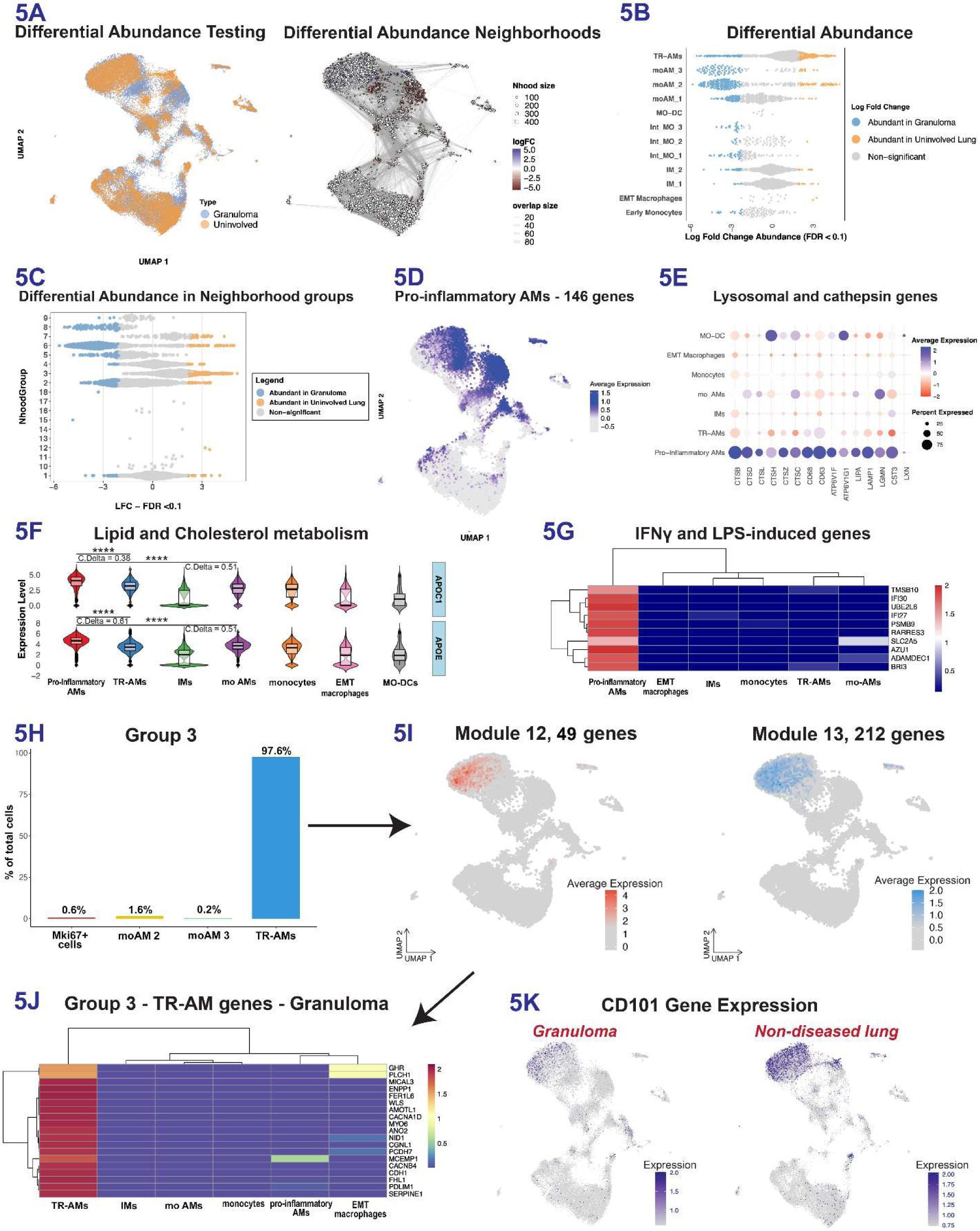
Identification of pro-inflammatory AMs in Granulomas. A. Differential abundance analysis of granuloma and uninvolved lung cells. (Left) UMAP visualization showing the distribution of cells from granuloma (blue) and uninvolved (yellow) lung tissue. (Right) Neighbors graph projected onto the UMAP plot, displaying cell cluster relationships based on differential abundance in granuloma and non-diseased lung populations. Neighbors with a significant Log2 Fold Change (LFC) increase in granuloma are shown in red, while those enriched in non-diseased lung are shown in blue (FDR < 0.1). B. Horizontal scatter plot showing changes in macrophage subset abundance, with color-coded data points indicating statistical significance and direction of change: Blue for neighbors with increased abundance in granuloma (FDR < 0.1), yellow for neighbors with higher abundance in uninvolved lung (FDR < 0.1), and light gray for no significant change. C. Horizontal scatter plot showing neighborhood (Nhood) clustering based on differential abundance, as calculated by changes in their LFC depletion rates. NhoodGroups are derived by aggregating neighbors (5A) that share similar transcriptional profiles and spatial proximity. NhoodGroups with increased abundance in granuloma are highlighted in blue, those with increased abundance in uninvolved lung are highlighted in yellow, and those with no significant change are shown in gray. D. Visualization of a granuloma-specific scWGCNA-derived module (146 genes) expression levels mapped onto UMAP embeddings. This module is specifically expressed by cells from NhoodGroup 6 and is exclusive to a subset of pro-inflammatory AMs in granuloma. E. DotPlot showing expression levels of lysosomal and cathepsin genes across various macrophage clusters. Dot size represents the percentage of cells expressing each gene, while color intensity indicates the average expression level. F. Violin plots showing the expression levels of lipid and cholesterol metabolism genes *Apoc1* and *Apoe* across various macrophage subsets. Each violin plot displays the distribution of expression levels, with overlaid box plots representing the median (center line), interquartile range (IQR; box bounds), and whiskers extending to 1.5 times the IQR. Cliff’s Delta (C. Delta) values indicate the effect size. Statistical significance is denoted by asterisks (**** = p < 0.0001). G. Heatmap showing the expression levels of IFNγ and LPS-induced genes across various macrophage subsets. Gene expression levels are color-coded, with red indicating higher expression and blue indicating lower expression. Hierarchical clustering is applied to both genes and cell populations to reveal patterns of co-expression and group similarities. H. Bar plot showing the distribution of cell types within Neighborhood Group 3. The plot displays the percentage of total cells contributed by each cell type, with TR-AMs comprising the majority (97.6%) of the group. I. Visualization of scWGCNA-derived gene module expression levels mapped onto UMAP embeddings. (Left) Expression pattern of genes in Module 12 (49 genes). (Right) Expression pattern of genes in Module 13 (212 genes). J. Heatmap displaying the expression levels of genes from Module 12 in granuloma across various macrophage subsets. These genes are specifically expressed by TR-AMs from Neighborhood Group 3. Gene expression levels are color-coded, with red indicating higher expression and blue indicating lower expression. Hierarchical clustering is applied to both genes and cell populations to reveal patterns of co-expression and group similarities. K. UMAP visualization of *Cd101* gene expression in granuloma (left) and non-diseased lung tissue (right). The expression levels are color-coded, with darker shades of blue indicating higher expression. The figure highlights the differential expression of *Cd101* between granuloma and non-diseased lung cells, with prominent expression in TR-AMs from non-diseased lung.

To elucidate the functional roles of these enriched populations, we performed scWGCNA on the 26,479 cells isolated from granulomas, identifying a 146-gene module unique to group 6 macrophages (Figure 5D). This module exhibited a pro-inflammatory signature, encompassing genes involved in lysosomal and phagolysosomal functions (e.g., CTSB, CTSD, CTSL), antioxidant and iron metabolism (GPX1, ATOX1, FTL, FTH1, GSTP1), lipid metabolism and cholesterol efflux (APOC1, APOE), complement activation (C1QA, C1QB, C1QC, CFD, CD59), and IFN-γ/LPS responses (GLUT5, ADAMDEC1, AZU1, RARRES3, TMSB10, BRI3) (Figure 5E–G, Supplementary Figures 6B–C, Supplementary Data 6)[84–95]. This pro-inflammatory profile suggests that both TR-AM and moAM subsets in granulomas actively contribute to the inflammatory milieu. Trajectory and pseudotime analyses supported these findings by showing that group 6 cells converge on terminal fates corresponding to TR-AMs (fate 13) and moAMs (fates 9, 18, 20, 24) (Figure 2G).

We then focused on subpopulations reduced of lesser abundance in granulomas. Group 3 emerged as a distinct subset comprising only TR-AMs (Figure 5H). Two gene modules were associated with this group (Figure 5I). Module 12 (49 genes) included factors that suppress NF-κB–mediated inflammation (PDLIM1, ENPP1, FHL1, NID1, SERPINE1, MCEMP1, GHR, WLS), calcium-activated enzymes (FER1L6, CACNB4, CACNA1D, ANO2), and mediators of actin-dependent processes and cell-cell interactions (LIMCH1, PCDH7, MYO6, MICAL3, AMOTL1, CDH1, CGNL1) (Figure 5J; Supplementary Data 7)[96–113]. In contrast, module 13 (212 genes) reflected the TR-AM–typical transcriptional signature found in non-diseased lung tissue, including PPARG, MRC1, PCCA, CD36, ABCG1, and LPL (Figure 4F, Supplementary Figure 5G, Supplementary Data 8). Expression of CD101, a marker linked to regulation of T-cell activity via reduction of IL12RA and IL-2 secretion[114–117], was lower in granulomas and present only in group 3 TR-AMs (Figure 5I–K). This decrease likely corresponds with increased macrophage activation and reduced numbers of anti-inflammatory TR-AMs in granulomas.

In summary, we identified two distinct TR-AM populations within granulomas: (1) group 6 TR-AMs, enriched in granulomas and displaying pro-inflammatory features, and (2) group 3 TR-AMs, reduced in granulomas and associated with lower pro-inflammatory activity. Despite these functional differences, both populations lacked CD14 staining (Figure 2A) and showed embryonic origins, as indicated by the absence of expression of the monocytic transcriptional signature identified in Figure 4A. These observations align with murine TB models showing pre-existing TR-AMs endowed with epigenetically driven pro-inflammatory potential in the lung prior to infection [16].

### Interstitial macrophages increase expression of canonical pro-inflammatory signatures in granuloma and are located in proximity of the granuloma core

Focusing on IMs, we first characterized their transcriptional profiles in non-diseased lung tissue. As detailed previously, using scWGCNA, we identified two gene modules that were specifically expressed by this macrophage ontogeny (Figure 6A). The first module (39 genes) contained components essential for NLRP3 inflammasome activation and IL-1β signaling (Figure 6B; Supplementary Data 9). The second module (60 genes) reflected M1 macrophage polarization, harboring genes linked to LPS and TNF stimulation, such as TNFAIP2, POU2F2, IRAK2, RFTN1, SLC7A5, ACSL3, MX2, ARID5B, FOXO1, and IER3 (Figure 6B and Supp. Figure 6D; Supplementary Data 10)[74, 118–130]. Consistent with this pro-inflammatory profile, immunostaining confirmed the functional phenotype of these IMs, as evidenced by their expression of CD80 and CD1d (Figure 1G).

**Figure 6.**
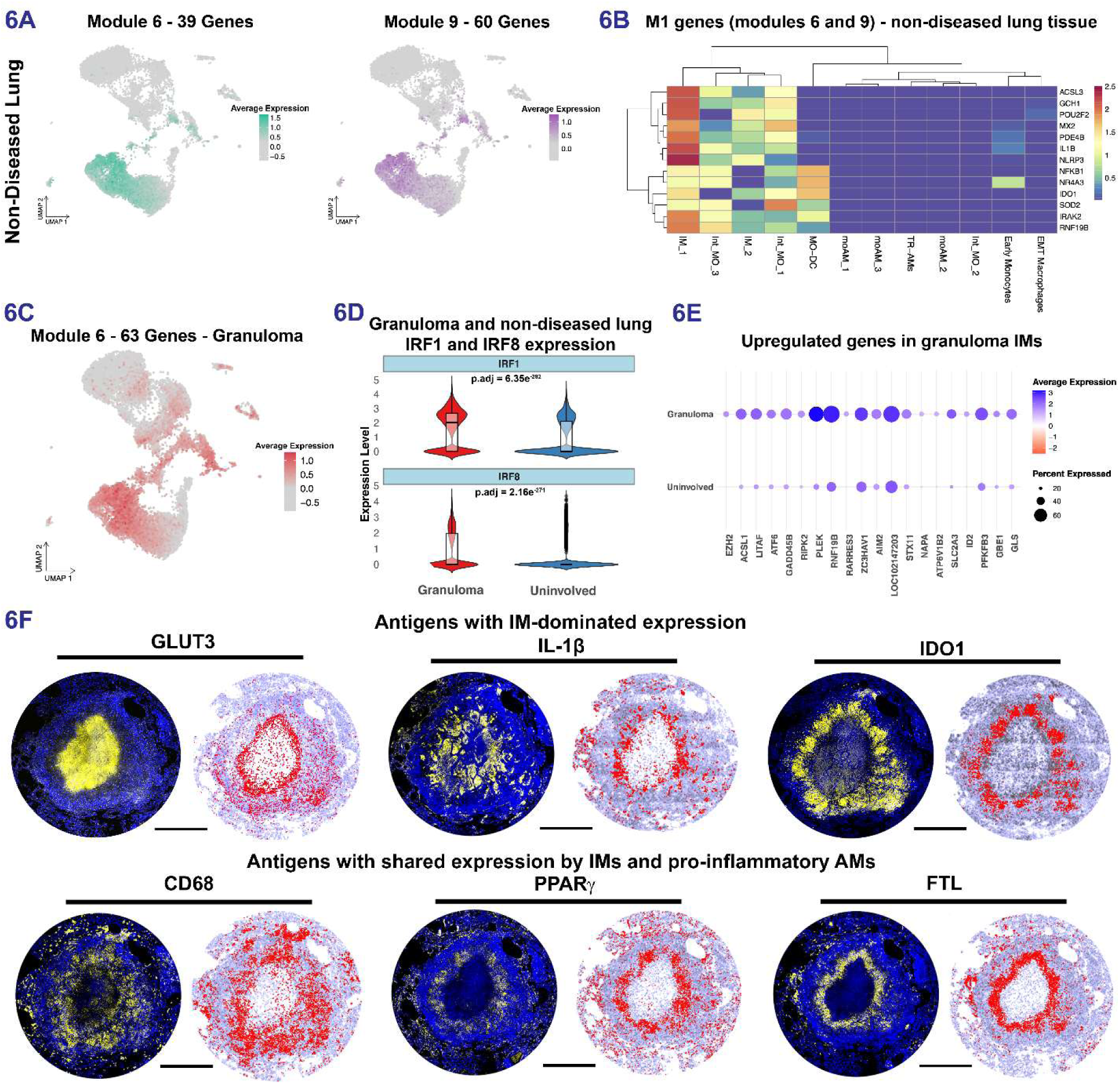
Analysis of IMs from both granuloma and non-diseased lung. A. Visualization of scWGCNA-derived gene module expression levels mapped onto UMAP embeddings across non-diseased lung cells. (Left) Expression pattern of genes in Module 6 (39 genes). (Right) Expression pattern of genes in Module 9 (60 genes). B. Heatmap displaying the expression levels of M1 genes from Modules 6 and 9 in non-diseased lung tissue across various macrophage subsets. Gene expression levels are color-coded, with red indicating higher expression and blue indicating lower expression. Hierarchical clustering is applied to both genes and cell populations to reveal patterns of co-expression and group similarities. C. Visualization of average expression levels of the scWGCNA-derived Module 6 (63 genes) in granuloma cells, mapped onto UMAP embeddings. D. Violin plots showing the expression levels of *Irf1* and *Irf8* in granuloma and non-diseased lung IMs. The expression levels are represented on the y-axis, with granuloma IMs shown in red and non-diseased lung IMs shown in blue. Each plot includes an overlaid box plot, where the center line indicates the median expression level, the box represents the interquartile range (IQR), and the whiskers extend to 1.5 times the IQR. Statistical significance between granuloma and non-diseased lung cells was assessed using a two-sided Wilcoxon Rank-Sum test, with p-values adjusted for multiple testing using the Benjamini-Hochberg method (p.adj). The adjusted p-values for the comparisons are displayed above each plot. E. Dot plot showing the expression levels of genes that are upregulated in IMs from granuloma compared to non-diseased (uninvolved) lung tissue. The size of the dots indicates the percentage of cells expressing each gene, while the color intensity represents the average expression level, with blue indicating upregulation. F. Localization of antigens that are strongly expressed by IMs (top row) and antigens expressed by both Ims and pro-inflammatory AMs (bottom row) in caseous granulomas. For each antigen, the panel on the left represents the microscopy image and the panel on the right is the plotted positions of antigen-positive cells (red) plotted against all of the nuclei for all of the cells in the image (blue). Scale bar = 500 μm.

When we restricted our analysis to IMs within granulomas, we discovered a 63-gene module that, akin to non-diseased lung IMs, showed robust IL-1β expression and included genes linked to M1 pro-inflammatory polarization (Figure 6C; Supplementary Data 11). A comparative analysis revealed that granuloma IMs showed increased expression of the macrophage regulome transcription factors IRF8 and IRF1. These transcription factors (Figure 6D), activated by IFN-γ in a STAT1-dependent manner, modulate key transcriptional programs central to inflammation and antimicrobial defense mechanisms against Mtb[131, 132]. Consequently, compared to non-diseased lung, granuloma IMs upregulated genes associated with pro-inflammatory cytokine production (e.g., EZH2, ACSL1, LITAF, ATF6, GADD45B, RIPK2) and phagosome/lysosome fusion and antimicrobial defense (e.g., PLEK, NKLAM, RARRES3, ZC3HAV1, AIM2, GBP1, STX11, NAPA, ATP6V1B2)[84, 133–146]. Additionally, they exhibited higher expression of genes related to glucose and glutamine/glutamate metabolism (e.g., GLUT3, ID2, PFKFB3, GBE1, GLS), consistent with the metabolic reprogramming towards glycolysis characteristic of M1 macrophages. This metabolic adaptation aligns with the critical role of glutamine/glutamate metabolism in promoting pro-inflammatory responses during TB[147, 148] (Figure 6E; Supplementary Data 12).

Immunostaining of granulomas confirmed the expression of IM-associated antigens, including GLUT3, IL-1β, and IDO1, in areas adjacent to the granuloma’s necrotic core (Figure 6F). Some IM antigens, such CD68, PPARG, and FTL (Figure 6F), were also expressed by pro-inflammatory AMs, suggesting these macrophages may undergo overlapping transcriptional programs during activation, despite residing in different microenvironments. In summary, IMs are abundant in granulomas and display specific pro-inflammatory profiles that distinguish them from IMs in non-diseased lung. Their antimicrobial phenotypes and metabolic reprogramming not only set them apart from alveolar macrophages but also suggest critical roles in mediating local immune responses against Mtb.

### Mtb reside in multiple macrophage subsets during early infection

We next sought to identify which host macrophage populations are preferentially infected by Mtb during the acute phase of the disease and elucidate how Mtb infection influences macrophage function. To visualize the bacteria *in situ*, we stained for mCherry (Mtb) and used CD11c as a general macrophage marker. To better characterize infected cells, we also stained for perilipin-2 (plin-2), a protein expressed by foamy macrophages and linked to caseation in human granulomas[149]. While mCherry^+^ Mtb were detected extracellularly in necrotic granuloma regions, consistent with previous findings in NHP granulomas [5], we also observed bacteria-rich cells at the granuloma’s outer edge (Figure 7A, regions 1, 2), where Mtb appeared as either singlets (region 1) or doublets (region 2) inside the cells. Phenotypically similar cells were also infected in non-caseous granulomas. Although plin-2^+^ macrophages were proximal to Mtb-infected cells (Figure 7A), mCherry^+^Mtb^+^ macrophages themselves did not express plin-2. These findings confirm the presence of both extracellular and intracellular Mtb within NHP granulomas and show that infection extends beyond the granuloma’s myeloid core during early TB. This could reflect early dissemination from the granuloma.

**Figure 7.**
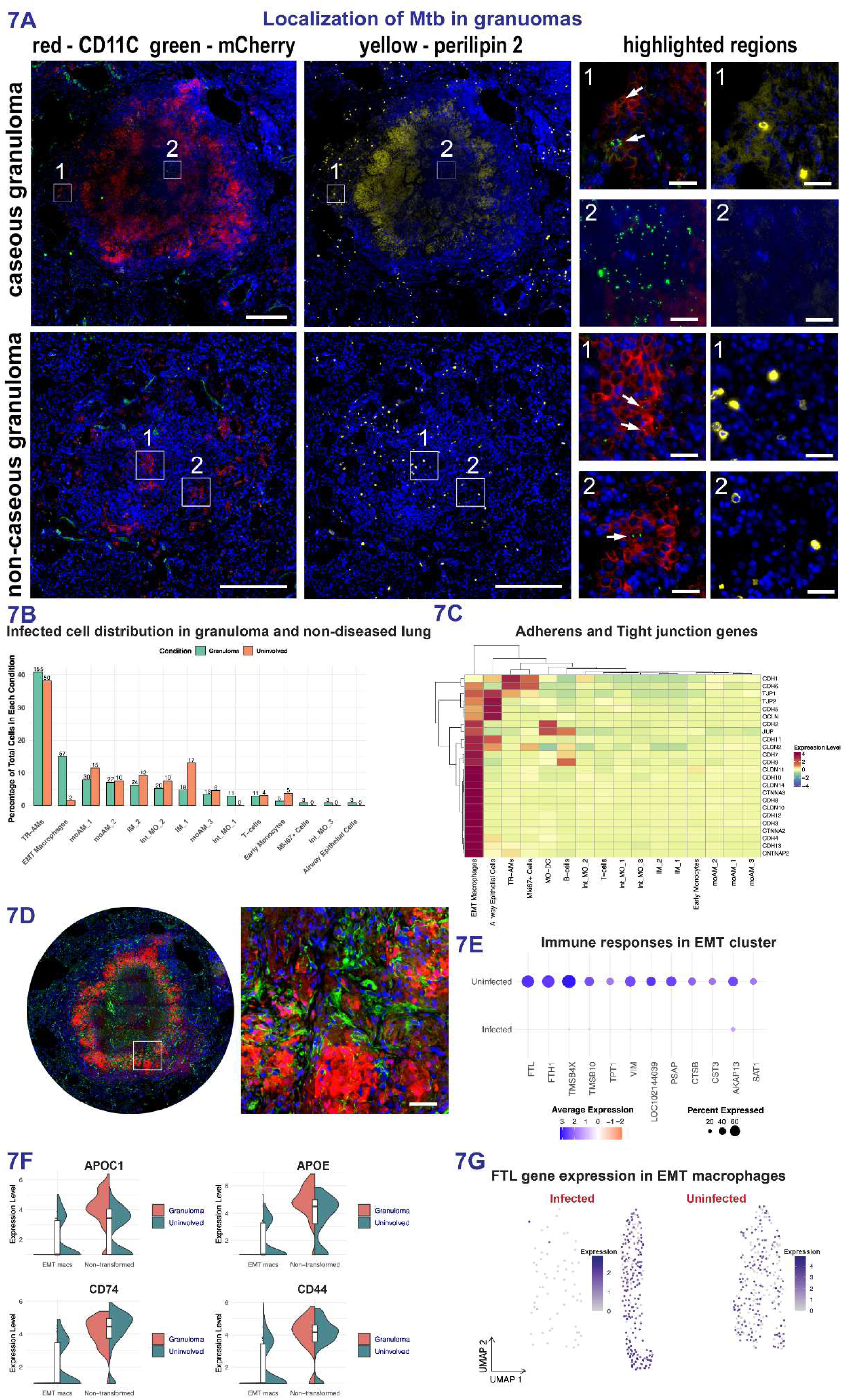
Distribution of Mtb-infected cells and identification of EMT macrophages. A. FFPE granulomas were stained for CD11c (red), mCherry (green), and perilipin-2 (plin-2, yellow) to visualize the presence of bacteria in the context of macrophages. Representative images of caseous and non-caseous granulomas are shown, with the position of Mtb in highlighted regions indicated by arrows. Enlarged views of the highlighted regions (labeled 1 and 2) show the presence of mCherry Mtb in necrotic (top) and non-necrotic (bottom) granulomas. Full granuloma scale bar = 500 μm, highlighted region scale bar = 50 μm. B. Bar plot showing the distribution of infected mCherry^+^cells across various cell types in granuloma and non-diseased lung tissue. The percentage of infected cells for each cell type, as a function of the total number of cells, is represented on the y-axis, with granuloma cells (green) and cells from non-diseased lung (orange). Cell subsets are listed along the x-axis with numbers above the bars indicating the raw count of infected cells recovered for each cell type within a category. C. Heatmap displaying the expression levels of adherens and tight junction genes across various cell types in granuloma and non-diseased lung tissue. Gene expression levels are color-coded according to the scale on the right, ranging from low (blue) to high (red). Hierarchical clustering is applied to both genes and cell populations to reveal patterns of co-expression and group similarities. D. Microscopy of IDO1 (red) and ITGB3 (green) expressing macrophages in the granuloma’s macrophage core. The highlighted region in the left panel is magnified in the right panel. Nuclei are stained with DAPI (blue). Inset scale bar = 50 μm. E. Dot plot showing the expression levels of immune response-related genes in the EMT macrophage cluster, comparing uninfected and infected cells. The size of the dots indicates the percentage of cells expressing each gene, while the color intensity represents the average expression level, with blue indicating higher expression and red indicating lower expression. F. Violin plots showing the expression levels of *Apoc1, Apoe, Cd74, and Cd44* in EMT macrophages (EMT macs) and non-transformed cells from granuloma and uninvolved lung tissue. The expression levels are represented on the y-axis, with granuloma cells shown in red and uninvolved cells shown in green. Each plot includes an overlaid box plot, where the center line indicates the median expression level, the box represents the interquartile range (IQR), and the whiskers extend to 1.5 times the IQR. The plots highlight the loss of macrophage marker expression in EMT macs specifically within granuloma tissue, while non-transformed cells maintain expression of these markers. G. UMAP plots showing expression of the *Ftl* gene in the EMT macrophage cluster in infected and uninfected samples. The left panel represents infected cells, while the right panel represents uninfected cells. Expression levels are color-coded, with darker shades of blue indicating higher expression levels. The plots highlight differences in *Ftl* gene expression between infected and uninfected EMT macrophages.

To further define the phenotype of Mtb-infected cells, we compared the transcriptional profiles of infected and uninfected cells that were flow-sorted based on their mCherry fluorescence. Because most bacteria reside extracellularly in the caseum, only a small number of mCherry^+^ cells were recovered by flow-sorting. After quality control, we retained n=511 mCherry^+^ cells, revealing that TR-AMs were the most frequently infected subset in both granulomatous tissue (155 of 380 infected cells) and tissue without gross evidence for disease (50 of 131 infected cells) (Figure 7B). Differential gene expression analysis of infected TR-AMs revealed upregulation of genes involved in negative modulation of innate inflammatory responses (ATG16L2, TFPI, SDC2, WLS, GPNMB, MARCH1, GDF15, CD9) and promotion of cell migration (ARHGEF28, MME, RAPH1, KDM3A, EXOC6B)[150–162] (Supp. Figure 7A), alongside downregulation of ribosomal proteins (Supplementary Figure 7B; Supplementary Data 13).

When comparing the infected cell subsets, we identified a unique cluster within granulomas that showed significantly higher levels of Mtb infection (Figure 7B). The cells comprising this population exhibited high expression of genes associated with adherens and tight junctions (e.g., cadherins, catenins, claudins, occludins), as well as adhesion- and polarity-related proteins (e.g., integrins, desmosomes), indicating a strong link to epithelial and mesenchymal phenotypes (Figure 7C). Their transcriptional profile paralleled cells undergoing epithelioid macrophage transformation (EMT macrophages), as described by Cronan et al. [163](Supplementary Data 14).

To validate this subset, we stained granulomas for IDO1, an IM-associated antigen, and ITGβ3, expressed by EMT macrophages in scRNA-seq. Immunostaining confirmed EMT macrophages as ITGβ3^+^, localized near the caseum-macrophage interface, interspersed with IDO1^+^ IMs (Figure 7D). scWGCNA analysis revealed a 66-gene module specific to EMT macrophages enriched for genes known to be associated with cytoskeletal organization, invasion, cell adhesion and positioning, and extracellular matrix (ECM) interactions (CTNNA3, PTPRD, SLIT3, EPHA6, CSMD1, SPOCK1, NRXN3, CNTN5, NAV2, CDH13, ERBB4, GRID2)[164–174] (Supplementary Data 15). Macrophage mesenchymal migration only occurs in pathological conditions, where tissue destruction is supported by specialized cell structures called podosomes that facilitate infiltration into pathological tissues[175–179]. We hypothesize that a similar transcriptional profile, enriched for pathways involving cytoskeletal organization, invasion and mesenchymal migration, would enable EMT macrophages to adopt an invasive mesenchymal-like state that facilitates their movement through TB-damaged lung tissue. Supporting this hypothesis, EMT macrophages expressed markers characteristic of mesenchymal and stromal cells (Supp. Figure 7C) and exhibited elongated morphologies, that differentiated them from the non-EMT macrophages in this granuloma region (Figure 7D, right). Moreover, EMT macrophages co-expressed both classical macrophage and mesenchymal markers, suggesting the presence of sub-structure within this population. To investigate further, we isolated and re-clustered the transcriptomes of EMT macrophages and identified substructure in this population that separated non-transformed macrophages within this subset and macrophages strongly characterized by their mesenchymal/stromal transformation. The latter subset was characterized by upregulated expression of EMT-associated genes and a concurrent loss of classical macrophage markers (Supp. Figure 7D and 7E). This phenotype was restricted to granulomas, and we found 57 of the 128 EMT cells recovered from granulomas were infected with Mtb (Figure 7B, Supp. Figure 7F).

Comparative transcriptional analysis between Mtb-infected and uninfected EMT macrophages revealed significant downregulation of genes critical for macrophage infection responses in infected cells, including ferritins (FTL, FTH1), actin sequestration (TMSB4X, TMSB10), and microtubule-stabilizing factors (TPT1) (Figure 7E-G, Supp. Figure 7G)[180–183]. In addition, expression of VIM, which promotes inflammatory cytokine release and oxLDL-induced TNF and IL-6 secretion, was abolished in infected EMT macrophages[184]. Antigen presentation pathways were similarly downregulated as evidenced by MHCII (LOC102144039) and lysosomal enzyme saposin (PSAP) — essential for presenting lipid antigens to CD1d-restricted T-cells[185] — and cathepsins (CTSB). This transcriptional changes extended to IFN-γ-mediated responses, with reduced expression of CST3, which is critical for this pathway[186]. Other key genes, including AKAP13, an activator of NF-kB downstream of TLR2[187], and SAT1, which plays a role in ferroptosis[188] were also downregulated (Supplementary Data 16). These data identify a novel pathway either induced or exploited by Mtb that would favor its survival and growth. Furthermore, EMT macrophages demonstrated reduced expression of lipid-metabolizing enzymes (APOC1, APOE), the MIF receptor (CD74), and the ECM receptor (CD44) (Figure 7F). Altogether, these findings suggest that EMT macrophages might be metabolically and transcriptionally reprogrammed to create an environment permissive to Mtb.

Taken together, our results indicate that TR-AMs are the dominant infected subset in early TB, exhibiting muted immune responses. In addition, a granuloma-specific EMT process produces a unique macrophage population that appears highly susceptible to Mtb infection. The extensive transcriptional alterations in EMT macrophages, including the suppression of crucial antimicrobial pathways, suggest they may serve as key players in disease progression and represent potential targets for therapeutic intervention.

## Discussion

Macrophages play key roles in TB pathogenesis, yet their phenotypic and functional diversity within granulomas, and their interactions with Mtb *in situ,* are not well understood. We addressed this knowledge gap by combining multimodal scRNA sequencing with spatial visualization to identify novel aspects of macrophage biology in granulomas from Mtb-infected NHPs, a model that closely mimics human TB pathogenesis. We uncovered a diverse macrophage landscape within granulomas, comprising subsets of TR-AMs, moAMs and IMs in different polarization and differentiation states. We also identified TR-AMs and a novel subset of EMT macrophages as the cell populations most permissive to *Mtb* infection during the early stage of infection. These infected macrophages exhibited blunted antimicrobial responses, aligning with murine studies that suggest *Mtb* preferentially infects and replicates in permissive cell subsets in early infection[12, 17]. The enrichment of these cells in early granuloma formation underscores the importance of understanding early host-pathogen interactions that may influence disease progression.

The interaction between macrophages and Mtb have been difficult to assess in humans and much of our current understanding comes from *in vitro* studies, murine TB studies, and research on resected tissue from patients with severe or chronic TB. This approach has made it difficult to investigate key aspects of TB progression in humans including identification of which cells are infected within granulomas, where these cells are located within the granuloma’s spatial context, and how Mtb influences the functional state of these cells. Moreover, the early events in tissues prior to development of adaptive immunity are even less understood but could have significant implications for later stages of disease progression. To investigate those questions in early TB, we used an NHP model with human-like pathophysiology and infection with mCherry-expressing Mtb to visualize the bacteria *in situ* and to phenotype the Mtb-infected cells in granulomas. We found extracellular bacteria in caseum, as previously noted [5], and intracellular bacteria in both caseous and non-caseous granulomas. Further phenotyping of these cells revealed multiple subsets of Mtb-infected cells but also highlighted an apparent preference for TR-AMs as host cells. This finding is consistent with previous work in mice where TR-AMs were permissive for Mtb [17, 189] and suggests that even after acute Mtb infection, TR-AMs may play unappreciated roles in bacterial persistence, proliferation, and dissemination.

One of the more striking revelations of our work is the identification and characterization of an EMT-like process within a subset of granuloma macrophages, a phenomenon widely studied in cancer that has received little investigation in TB. Intriguingly, we found a high proportion of the Mtb-infected macrophages were undergoing EMT, which was characterized by increased expression of genes involved in cell adhesion, cytoskeletal remodeling, and extracellular matrix interactions. A further analysis comparing the infected to uninfected cells within this subset suggests that Mtb may be actively undermining macrophage defenses by downregulating immune functions. Alternatively, we may be detecting cells with preexisting immune deficits in which Mtb can persist and proliferate. Future work investigating this relationship, including at different stages of TB, will yield information on macrophage-centric factors the correlate with bacterial permissiveness in granulomas.

Our study provides a foundation for future research identifying the molecular mechanisms driving differential macrophage susceptibility to Mtb infection. Understanding the implications of these transcriptional alterations on macrophage functionality and the overall immune response to Mtb infection is needed because macrophage behavior directly influences bacterial containment, granuloma formation, and progression to active disease. Early alterations that impair macrophage antimicrobial functions or promote a permissive environment for Mtb can shape long-term infection outcomes and limit the efficacy of host-directed therapies[190]. Expanding analyses to later infection stages and utilizing genetic or pharmacological interventions to modulate pathways identified in macrophages will be critical for verifying their contribution to Mtb pathogenesis and assessing their viability as targets for improving host resistance to Mtb infection.

Our study has limitations that need to be considered in the interpretation of our data. First, we focused on the early stage of disease in a NHP species that is before development of robust adaptive immunity and is permissive for Mtb replication. Considering that, we are examining granulomas prior to their segregation into protective and non-protective lesions[190]; thus, we were not able to define macrophage subsets and lesion-level correlates of immunity. Future experiments investigating stages of the disease where granulomas are more restrictive for Mtb replication may improve our understanding of how granuloma macrophage function and population composition changes as immunity matures in TB. Second, our sample size was limited, and future studies with larger cohorts across different disease stages will provide a more comprehensive understanding of granuloma macrophage dynamics. Finally, while we identified novel transcriptional and spatial features of macrophage subsets, functional validation is necessary to confirm their roles in TB progression. likewise, we lacked the ability to determine if the differences we noted between macrophage infection status and immune function were based on cause-effect relationships driven by the bacterium or the Mtb-infected cells had pre-existing deficits in their antimycobacterial immune capacity that allowed us to detect bacteria in these cells. Future studies investigating macrophage responses *ex vivo* may provide insight into aspect of host-pathogen interaction in TB.

In conclusion, our study advances the understanding of the macrophage ecosystem in TB granulomas, revealing the phenotypic complexity of macrophage subsets during Mtb infection in NHPs and identifying potential processes that Mtb exploits to improve its survival. This research provides a basis for future work aimed at developing targeted interventions against TB. The diversity in macrophage responses, especially the distinct functions of moAMs and TR-AMs, emphasizes the importance of nuanced therapeutic strategies. These strategies should account for the different roles and states of macrophages in the Mtb-infected lung.

## Acknowledgements

The Authors acknowledge Jamie Tomko, Carolyn Bigbee, and the Flynn Lab technician and veterinary teams for their contributions to the NHP studies. This work was funded by NIH AI134183 to DGR and JLF, and OD032135 to DGR. Support for JTM was provided in part by NIH grants AI167710 and AI164970. Additional support was provided in part by funding by the Wellcome Leap ΔTissue Program awarded to (JLF and JTM). We gratefully acknowledge NIH UC7AI180311 from the National Institute of Allergy and Infectious Diseases for supporting the Operations of The University of Pittsburgh Regional Biocontainment Laboratory within the Center for Vaccine Research.

## Author contributions

Conceptualization: DGR, JLF, JTM, DP; Methodology: JTM, DP; Software: DP, Validation: DGR, JLF, JTM, DP; Formal analysis: DP, JTM; Investigation: PSS, MLN, BFJ, MAR, PLL, HJB, PM, MAR, DP, JTM; Resources: DGR, JLF, JTM; Data curation: JTM, DP; Writing – original draft: JTM, DP; Writing – reviewing and editing: all authors; Visualization: SSP, MLN, HJB, JTM, DP; Supervision: DGR, JLF, JTM, DP, CAS; Project administration: JLF, DGR, CAS, JTM, DP; Funding acquisition: JLF, JTM, DGR.

---

**Supplementary Figure 1.**
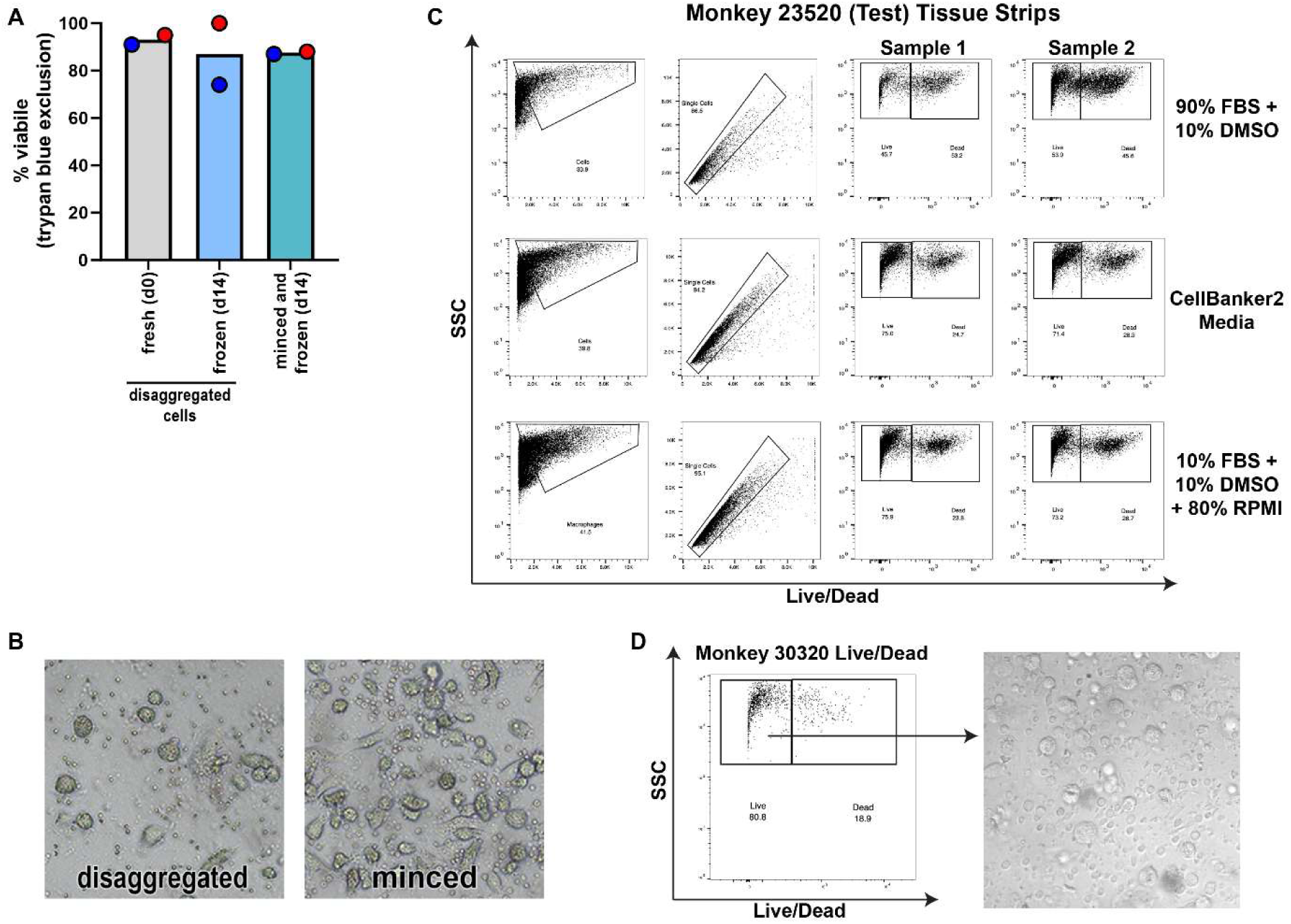
Cryopreserved lung tissue yields viable cells after being thawed. A. The viability of freshly disaggregated cells was compared against tissues that were disaggregated and then frozen, or tissues that were minced before freezing and then disaggregated after being thawed were compared. The frozen samples were stored at -80C for 14 days before being thawed and processed for culturing cells or for flow cytometry. Each color represents a different monkey and the bar represents the mean value of non-diseased lung samples from two animals. B. Phase contrast microscopy of cells from cryopreserved tissues were thawed after 14 days at -80C and cultured in RPMI+10% FBS before being imaged by to show populations of viable alveolar macrophages (large granular cells) amidst smaller lymphocyte-sized cells. C. Flow cytometric comparison of the viability of cells frozen with different cryopreservatives and shipped between facilities. D. Flow cytometry of and appearance of cells from minced tissue.

**Supplemental Figure 2.**
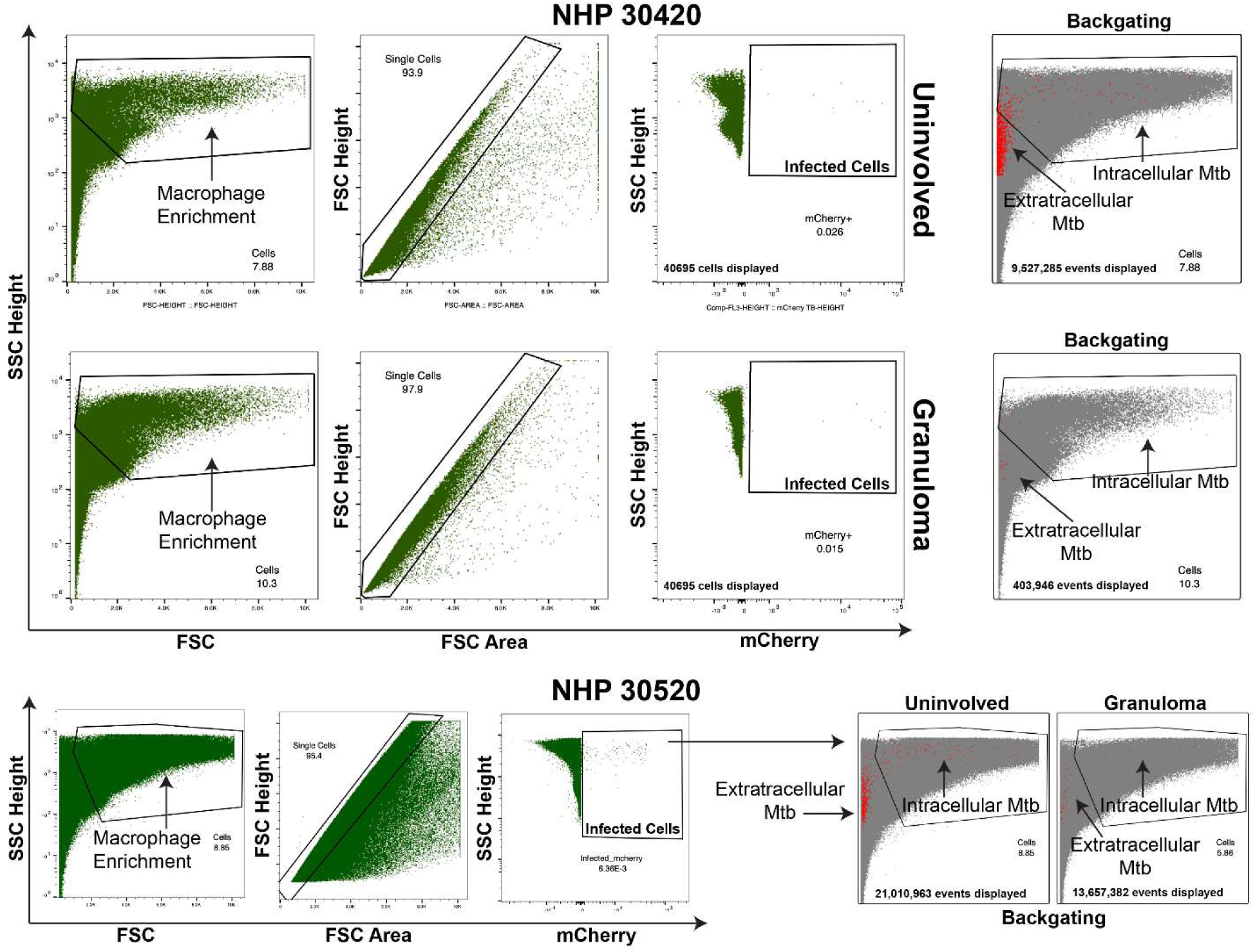
Gating strategy for sorting macrophages from NHP lung tissue including detection of Mtb-infected cells.

**Supplementary Figure 3.**
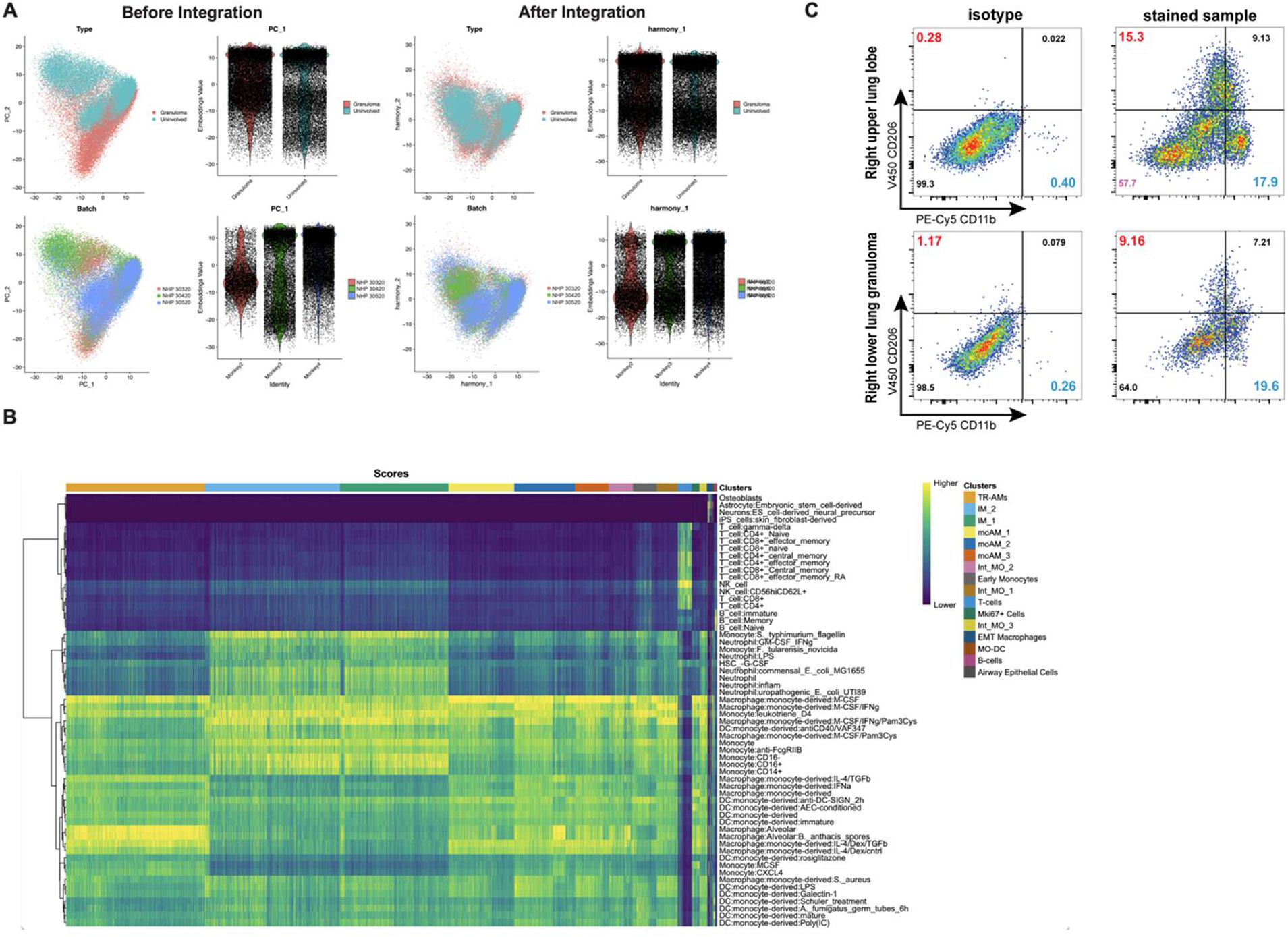
Alveolar and interstitial macrophages can be differentiated by CD206 and CD11b staining. A. PCA and violin plots visualizing the merged NHP datasets before and after integration with Harmony for the two covariates: type and batch. As visualized in the figure, data integration with Harmony allowed us to correctly identify shared identities from cells of different batches and tissue types. B. Heatmaps showing the results of reference-based analysis for all cells in the integrated dataset against the Human Primary Cell Atlas database. Higher similarity scores among the transcriptional profiles of query and reference datasets are showed in yellow. C. Lung tissue (top row) and lung granulomas (bottom row) were homogenized and stained for CD206 and CD11b and compared against isotype controls.

**Supplementary Figure 4.**
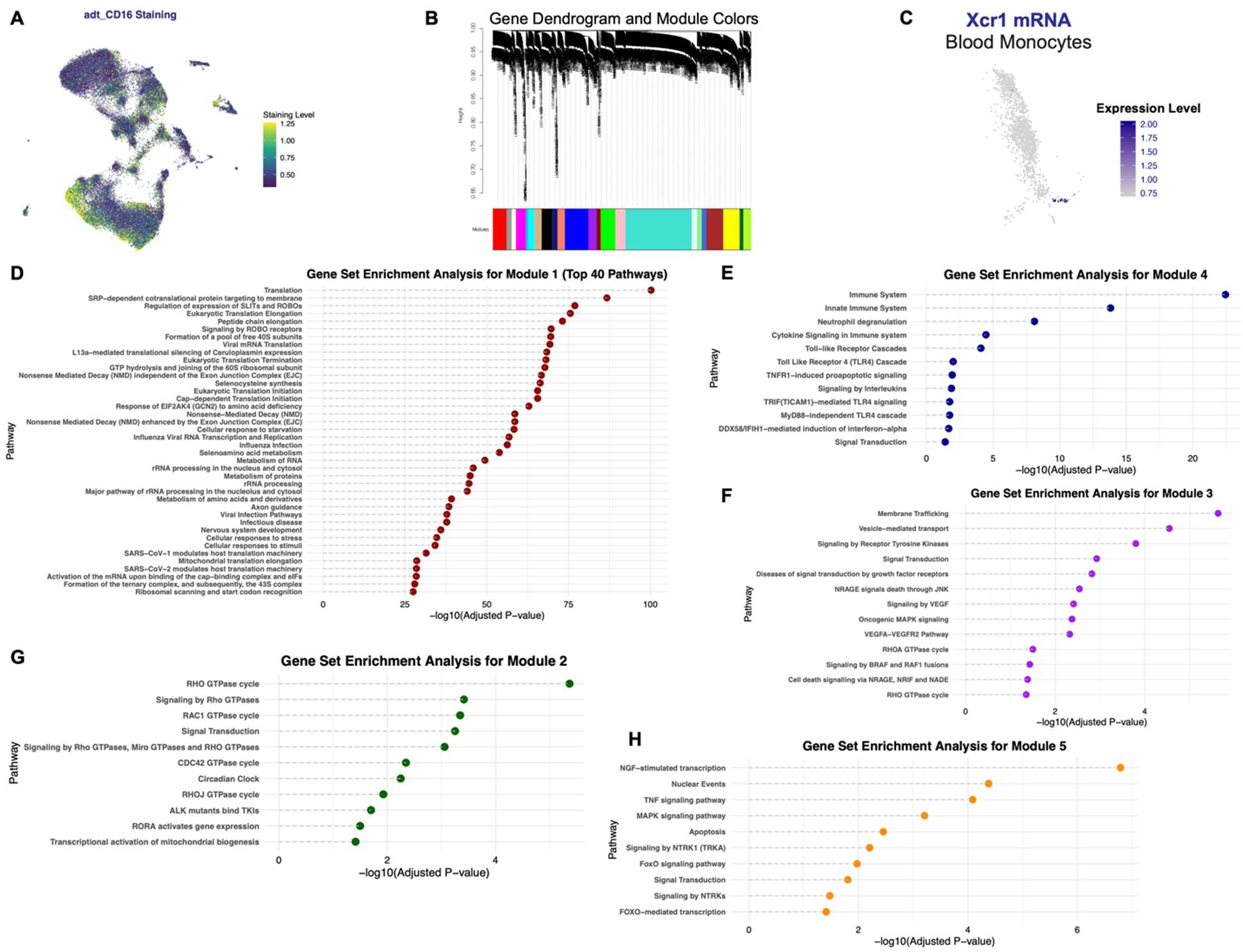
Trajectory and pseudotime analysis of monocyte-derived cells reveal multiple polarization programs in NHP-infected Mtb lungs. A. UMAP Visualization of CD16 Staining levels. This plot depicts the staining level for CD16 across individual cells projected onto a UMAP embedding. Each point represents a single cell, with the color gradient (from dark blue to yellow) indicating the relative staining level. B. Gene Dendrogram with 23 Co-expression Modules Generated by scWGNA. This dendrogram illustrates the hierarchical clustering of genes based on their expression profiles as analyzed by scWGNA. Twenty-three distinct gene modules were identified, with each branch of the dendrogram representing a cluster of co-expressed genes. These modules capture groups of genes that share similar expression patterns and may be associated with specific biological processes or cellular phenotypes. C. UMAP plot showing RNA expression levels of XCR1 in early monocytes. Cells are colored on a gradient from light grey (low expression) to dark blue (high expression) D. Gene Set Enrichment Analysis for Module 1. Dot plot displaying the top 40 enriched pathways for Module 1, with the x-axis representing –log₁₀(adjusted p-values) and the y-axis listing pathways ordered by significance. Dashed lines extend from the x-axis to each point, highlighting the degree of enrichment—higher values indicate stronger statistical significance. The analysis was conducted using g:Profiler with standard parameters. E. Gene Set Enrichment Analysis for Module 4. Dot plot showing enriched immune-related pathways in Module 4. The x-axis displays –log₁₀(adjusted p-values), reflecting the significance of enrichment, while the y-axis lists pathways ordered by their enrichment score. Dashed lines extend from each point to the x-axis to emphasize the magnitude of enrichment. The analysis was conducted using g:Profiler with standard parameters. F. Gene Set Enrichment Analysis for Module 3. Dot plot displaying enriched pathways for Module 3. Each purple point represents a pathway, with its position on the x-axis indicating the –log₁₀(adjusted p-value) and the dashed line illustrating its magnitude of enrichment. Pathways are ordered by significance, with higher –log₁₀ values reflecting stronger enrichment. The analysis was conducted using g:Profiler with standard parameters. G. Gene Set Enrichment Analysis for Module 2. Dot plot displaying enriched pathways for Module 2. Each dark green point represents a pathway, with its x-axis position reflecting the –log₁₀(adjusted p-value). Dashed lines extend from the x-axis to each point, illustrating the magnitude of enrichment—higher values indicate stronger significance. The analysis was conducted using g:Profiler with standard parameters. H. Gene Set Enrichment Analysis for Module 5. Dot plot displaying enriched pathways for Module 5, with the x-axis showing –log₁₀(adjusted p-values). Each orange point represents a pathway, and dashed lines extend from the x-axis to the points, illustrating the magnitude of enrichment (with higher values indicating stronger significance). The analysis was conducted using g:Profiler with standard parameters.

**Supplementary Figure 5.**
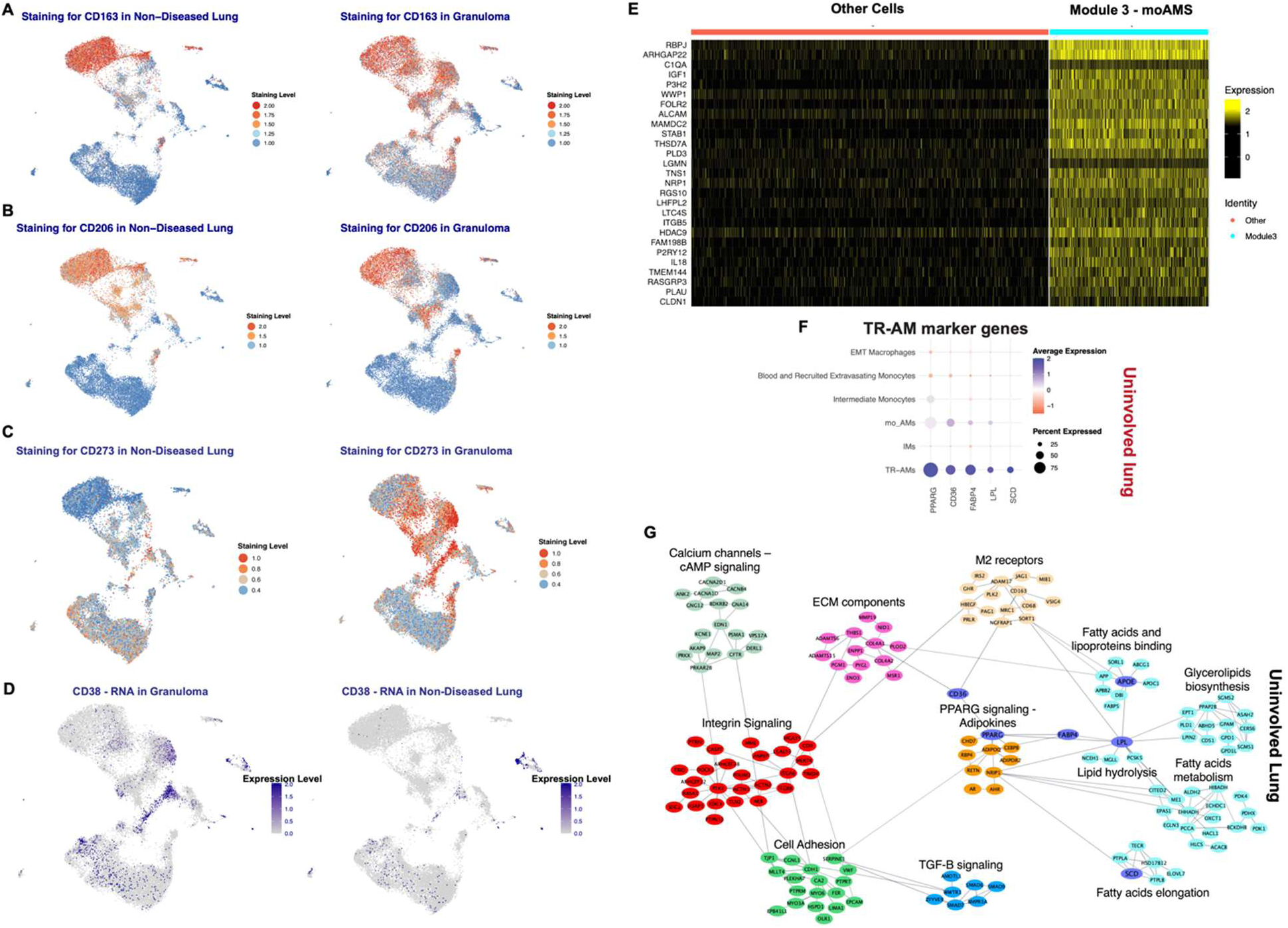
Antibody staining and gene expression analysis for different subsets of AMs. A. UMAP visualization of CD163 staining in Non-Diseased Lung and Granuloma. The plot displays CD163 staining levels (color-coded from blue for low to dark red for high expression) across cells from non-diseased lung and granuloma samples. B. UMAP visualization of CD206 staining in Non-Diseased Lung and Granuloma. The plot displays CD206 staining levels (color-coded from blue for low to dark red for high expression) across cells from non-diseased lung and granuloma samples. C. UMAP visualization of CD273 staining in Non-Diseased Lung and Granuloma. The plot displays CD273 staining levels (color-coded from blue for low to dark red for high expression) across cells from non-diseased lung and granuloma samples. D. UMAP visualization of CD38 RNA expression levels in Granuloma and Non-Diseased Lung. CD38 RNA levels are displayed using a color gradient from light grey (low expression) to dark blue (high expression). E. Heatmap showing the expression of genes linked to monocyte-to-macrophage differentiation in cells from uninvolved lung (downsampled to 6000 cells). The heatmap is split to compare moAMs with all other cells, with expression values scaled between –2.5 and 2.5. The color gradient—from black (low expression) to yellow (high expression)—illustrates relative gene expression levels. F. Dot plot showing the expression levels of TR-AM marker genes in uninvolved lung tissue. The size of the dots indicates the percentage of cells expressing each gene, while the color intensity represents the average expression level, with blue indicating upregulation. G. Pathway and protein-protein interaction network analysis for the scWGCNA 869-gene module that defines TR-AMs in non-diseased (uninvolved) lung tissue.

**Supplementary Figure 6.**
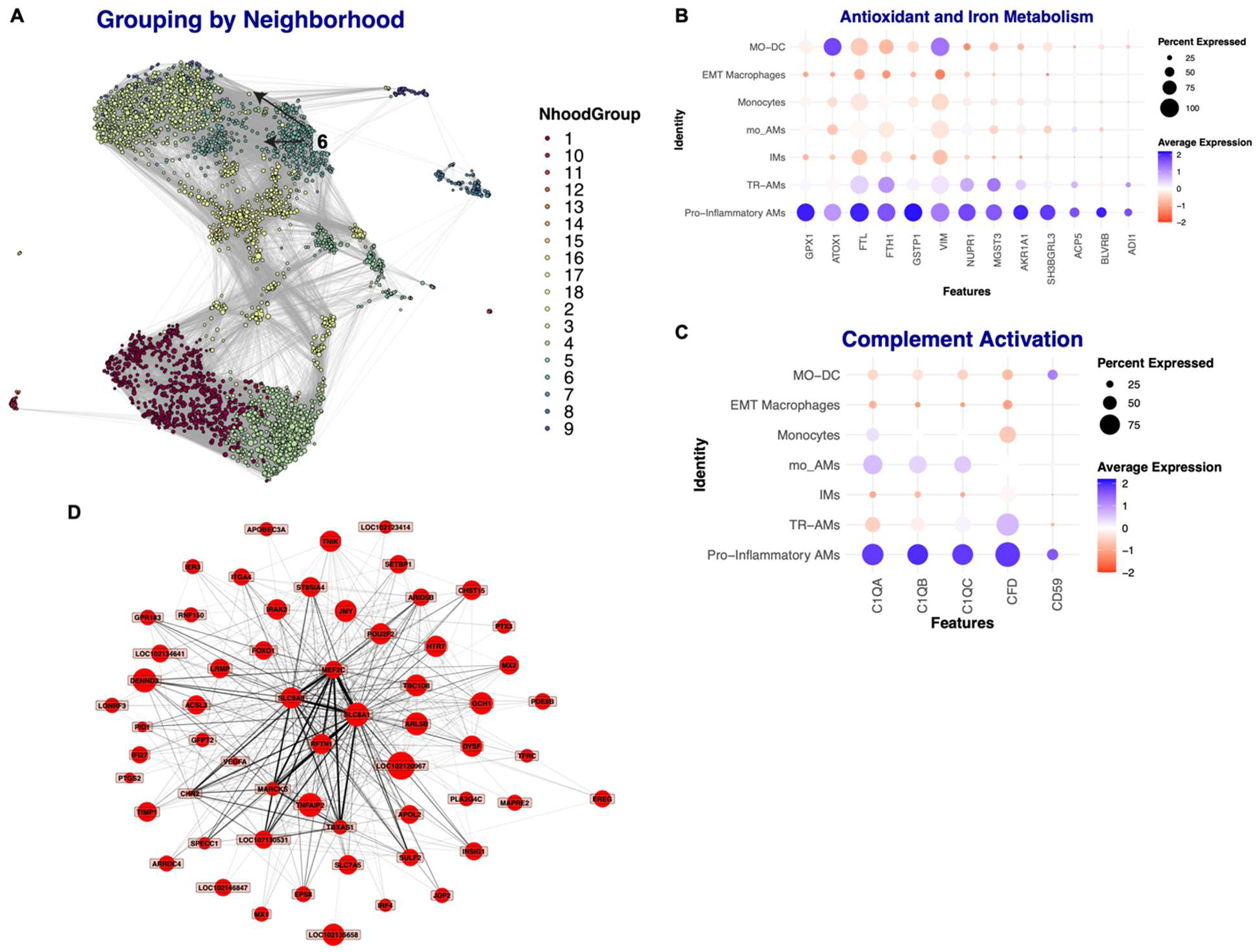
TR-AM and moAM contribute to the local pro-inflammatory milieu. A. UMAP visualization of 18 neighborhood groups identified by differential abundance analysis. Group 6 is highlighted with arrows for easier visual identification, and the UMAP layout represents the spatial distribution of these groups. B. Dot plot showing antioxidant and iron metabolism gene expression in pro-inflammatory AMs (Group 6 Nhood) in granuloma. Each dot reflects the percentage of cells expressing the gene (dot size) and the average expression level (color intensity), highlighting upregulation of antioxidant and iron metabolism pathways. C. Dot plot showing complement activation gene expression in pro-inflammatory AMs (Group 6 Nhood) in granuloma. Dot size indicates the proportion of cells expressing each gene, while color intensity represents the average expression level. D. scWGCNA network map for module 9 (60 genes) that defines IMs in non-diseased lung. The network displays co-expression relationships among genes enriched in response to LPS and TNF stimulation, including TNFAIP2, POU2F2, IRAK2, RFTN1, SLC7A5, ACSL3, MX2, ARID5B, FOXO1, and IER3. Each node represents a gene, and edges indicate the strength of co-expression as determined by the scWGCNA analysis.

**Supplemental Figure 7.**
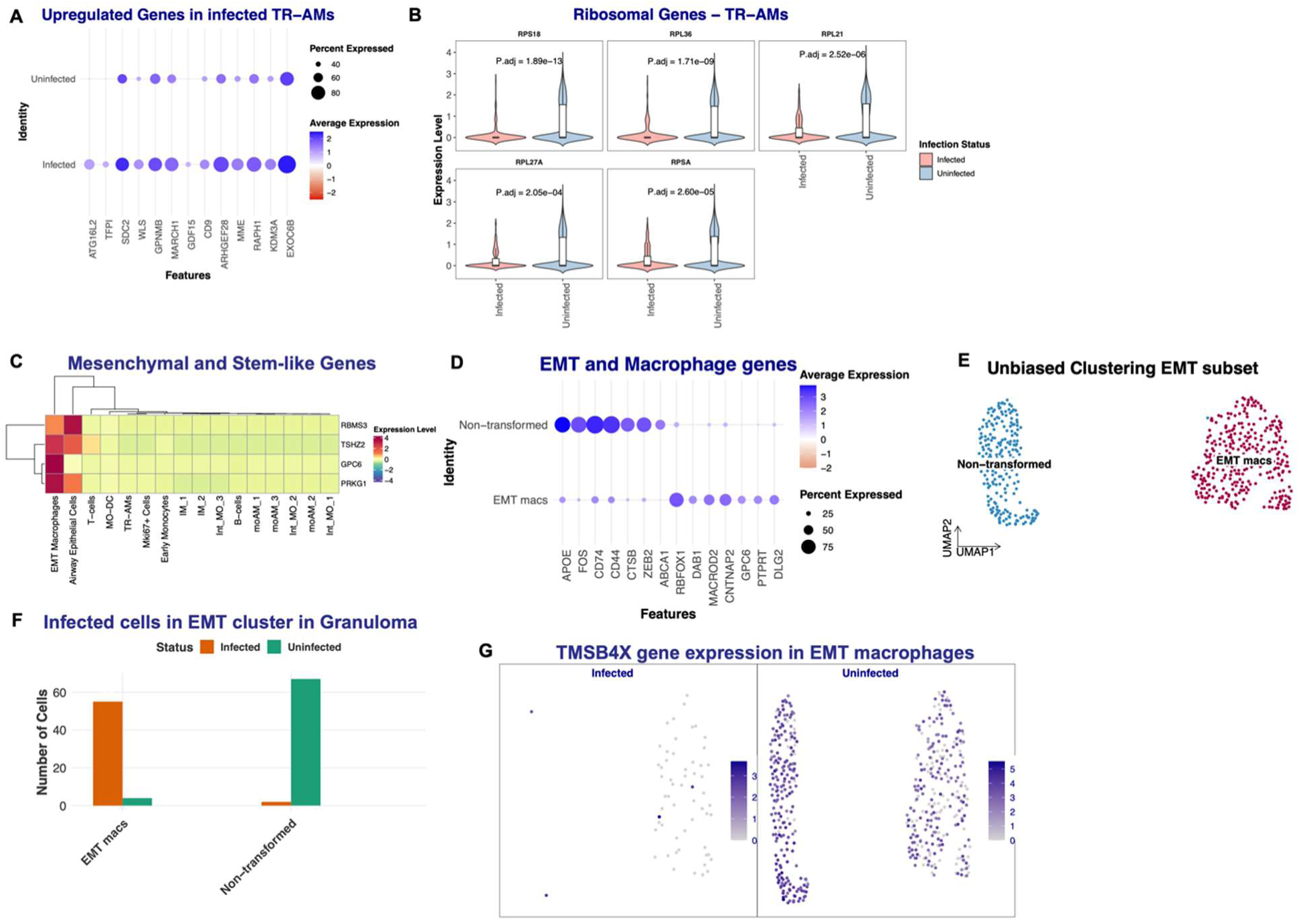
Infected and uninfected TR-AMs, and EMT and non-EMT macrophages differ. A. Upregulated genes in infected TR-AMs. The plot displays expression of 13 genes associated with negative regulation of innate inflammation and promotion of cell migration, in infected and uninfected TR-AMs. Dot size reflects the percentage of cells expressing each gene, and the color scale (red to blue, with white at 0; values limited between –2.5 and 2.5) indicates the average expression level. B. Violin plots of ribosomal gene expression in TR-AMs. The figure displays the expression distribution of five ribosomal genes (RPS18, RPL36, RPL21, RPL27A, RPSA) in TR-AMs, grouped by infection status. For each gene (faceted panel), violins illustrate the full expression distribution with overlaid boxplots summarizing the median and interquartile range. FDR-adjusted p-values indicate the statistical significance of expression differences between groups. C. Row-scaled expression for four mesenchymal and stem-like marker genes (RBMS3, TSHZ2, GPC6, and PRKG1) are shown. Expression values were averaged by cluster and visualized using a custom diverging color palette, with clustering based on correlation (genes) and Euclidean (clusters) distances. D. Dot plot showing expression of EMT and macrophage genes. The plot displays the expression levels of 14 genes associated with EMT and macrophage function. Dot size indicates the proportion of cells expressing each gene, while dot color reflects the average expression level on a diverging scale (red for lower, white for intermediate, blue for higher expression). E. Unbiased re-clustering of the EMT subset, lead to the identification of two separate populations called “Non-transformed” and “EMT macs”. F. Histogram visualization of the number of Mtb-infected cells in EMT Macs and Non-Transformed subsets. Counts for each cluster are displayed and colored by infection status (Infected vs. Uninfected), facilitating a direct comparison of infection prevalence between the two subsets. G. UMAP plots showing expression of the *Tmsb4x* gene in the EMT macrophage cluster among infected and uninfected cells. The left panel represents infected cells, while the right panel represents uninfected cells. Darker shades of blue indicate higher expression levels.

**Supplemental Table 1.**
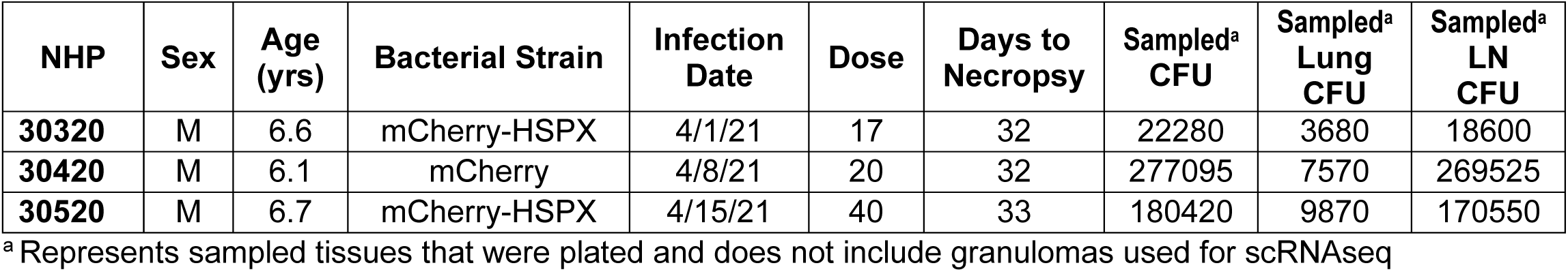

| **NHP** | **Sex** | **Age (yrs)** | **Bacterial Strain** | **Infection Date** | **Dose** | **Days to Necropsy** | **Sampled<sup>a</sup> CFU** | **Sampled<sup>a</sup> Lung CFU** | **Sampled<sup>a</sup> LN CFU** |
| --- | --- | --- | --- | --- | --- | --- | --- | --- | --- |
| **30320** | M | 6.6 | mCherry-HSPX | 4/1/21 | 17 | 32 | 22280 | 3680 | 18600 |
| **30420** | M | 6.1 | mCherry | 4/8/21 | 20 | 32 | 277095 | 7570 | 269525 |
| **30520** | M | 6.7 | mCherry-HSPX | 4/15/21 | 40 | 33 | 180420 | 9870 | 170550 |

<sup>a</sup> Represents sampled tissues that were plated and does not include granulomas used for scRNAseq

**Supplemental Table 2.**

| **NHP** | **Scan Date** | **Days Post Infection** | **Weeks Post Infection** | **Probe** |
| --- | --- | --- | --- | --- |
| **30320** | 4/30/21 | 29 | 4 | <sup>18</sup>F-FDG |
| **30420** | 5/6/21 | 28 | 4 | <sup>18</sup>F-FDG |
| **30520** | 5/14/21 | 30 | 4 | <sup>18</sup>F-FDG |

**Supplemental Table 3.**

| **NHP** | **Lymph node** | **CFU** |
| --- | --- | --- |
| **30320** | L Cran HLN1 | 460 |
| **30420** | LMS Br LN | 165200 |
|  | RMS Br LN | 9975 |
|  | L Carin LN | 18800 |
|  | R Carin LN | 66500 |
|  | L Cran HLN1 | 50 |
|  | L Cran HLN2 cran | 9000 |
| **30520** | R Subclav LN | 24500 |
|  | L Carin LN | 125500 |
|  | R Carin LN | 2075 |
|  | L Cran HLN1 | 6950 |
|  | L Cran HLN2 | 7225 |

**Supplemental Table 4.**
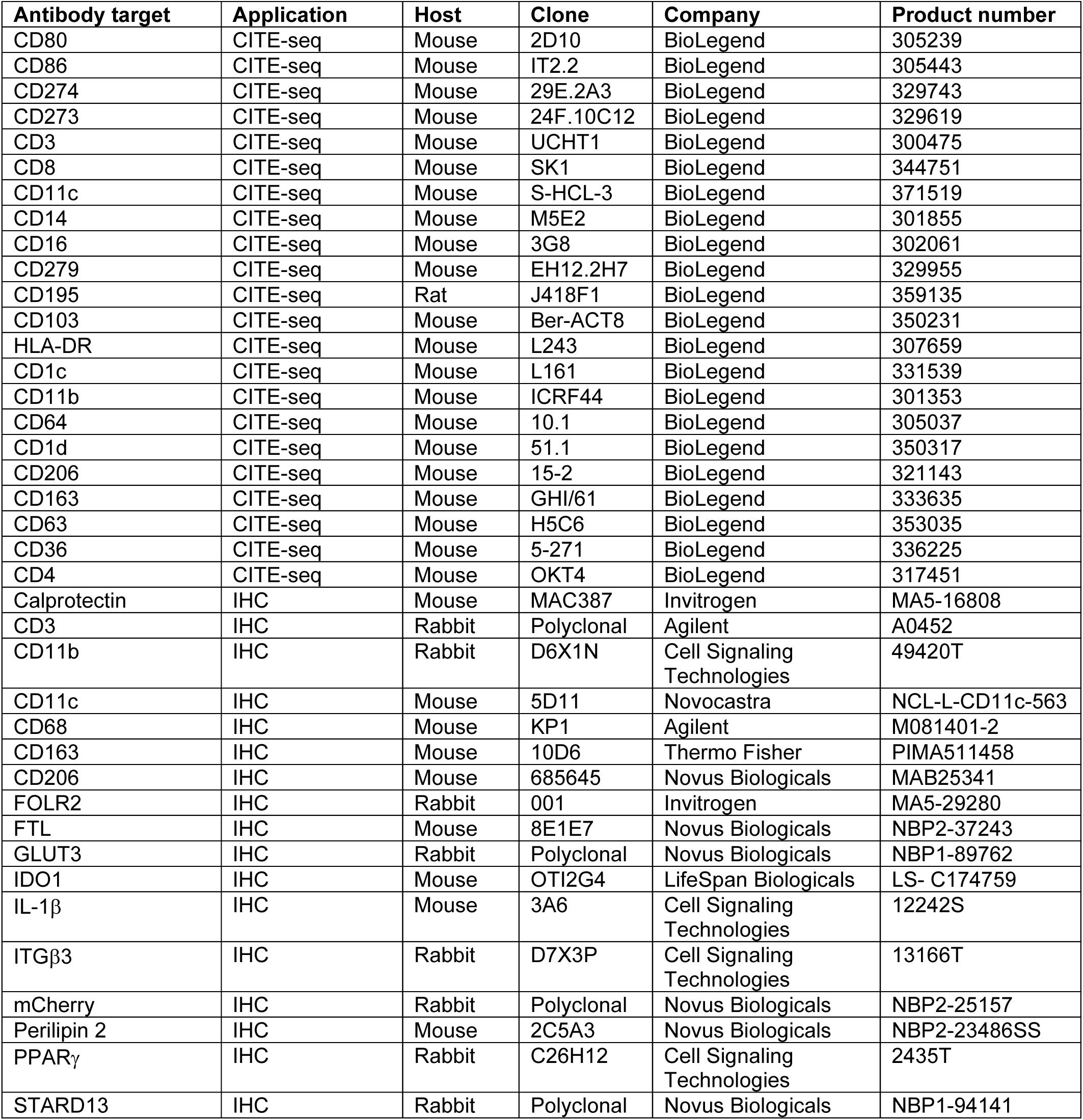
Antibodies used for CITE-seq and immunohistochemistry.

| **Antibody target** | **Application** | **Host** | **Clone** | **Company** | **Product number** |
| --- | --- | --- | --- | --- | --- |
| CD80 | CITE-seq | Mouse | 2D10 | BioLegend | 305239 |
| CD86 | CITE-seq | Mouse | IT2.2 | BioLegend | 305443 |
| CD274 | CITE-seq | Mouse | 29E.2A3 | BioLegend | 329743 |
| CD273 | CITE-seq | Mouse | 24F.10C12 | BioLegend | 329619 |
| CD3 | CITE-seq | Mouse | UCHT1 | BioLegend | 300475 |
| CD8 | CITE-seq | Mouse | SK1 | BioLegend | 344751 |
| CD11c | CITE-seq | Mouse | S-HCL-3 | BioLegend | 371519 |
| CD14 | CITE-seq | Mouse | M5E2 | BioLegend | 301855 |
| CD16 | CITE-seq | Mouse | 3G8 | BioLegend | 302061 |
| CD279 | CITE-seq | Mouse | EH12.2H7 | BioLegend | 329955 |
| CD195 | CITE-seq | Rat | J418F1 | BioLegend | 359135 |
| CD103 | CITE-seq | Mouse | Ber-ACT8 | BioLegend | 350231 |
| HLA-DR | CITE-seq | Mouse | L243 | BioLegend | 307659 |
| CD1c | CITE-seq | Mouse | L161 | BioLegend | 331539 |
| CD11b | CITE-seq | Mouse | ICRF44 | BioLegend | 301353 |
| CD64 | CITE-seq | Mouse | 10.1 | BioLegend | 305037 |
| CD1d | CITE-seq | Mouse | 51.1 | BioLegend | 350317 |
| CD206 | CITE-seq | Mouse | 15-2 | BioLegend | 321143 |
| CD163 | CITE-seq | Mouse | GHI/61 | BioLegend | 333635 |
| CD63 | CITE-seq | Mouse | H5C6 | BioLegend | 353035 |
| CD36 | CITE-seq | Mouse | 5-271 | BioLegend | 336225 |
| CD4 | CITE-seq | Mouse | OKT4 | BioLegend | 317451 |
| Calprotectin | IHC | Mouse | MAC387 | Invitrogen | MA5-16808 |
| CD3 | IHC | Rabbit | Polyclonal | Agilent | A0452 |
| CD11b | IHC | Rabbit | D6X1N | Cell Signaling Technologies | 49420T |
| CD11c | IHC | Mouse | 5D11 | Novocastra | NCL-L-CD11c-563 |
| CD68 | IHC | Mouse | KP1 | Agilent | M081401-2 |
| CD163 | IHC | Mouse | 10D6 | Thermo Fisher | PIMA511458 |
| CD206 | IHC | Mouse | 685645 | Novus Biologicals | MAB25341 |
| FOLR2 | IHC | Rabbit | 001 | Invitrogen | MA5-29280 |
| FTL | IHC | Mouse | 8E1E7 | Novus Biologicals | NBP2-37243 |
| GLUT3 | IHC | Rabbit | Polyclonal | Novus Biologicals | NBP1-89762 |
| IDO1 | IHC | Mouse | OTI2G4 | LifeSpan Biologicals | LS- C174759 |
| IL-1β | IHC | Mouse | 3A6 | Cell Signaling Technologies | 12242S |
| ITGβ3 | IHC | Rabbit | D7X3P | Cell Signaling Technologies | 13166T |
| mCherry | IHC | Rabbit | Polyclonal | Novus Biologicals | NBP2-25157 |
| Perilipin 2 | IHC | Mouse | 2C5A3 | Novus Biologicals | NBP2-23486SS |
| PPARγ | IHC | Rabbit | C26H12 | Cell Signaling Technologies | 2435T |
| STARD13 | IHC | Rabbit | Polyclonal | Novus Biologicals | NBP1-94141 |

